# Glial cells in the heart? Replicating the diversity of the myocardium with low-cost 3D models

**DOI:** 10.1101/306233

**Authors:** Jonathan R. Soucy, Jody Askaryan, David Diaz, Abigail N. Koppes, Nasim Annabi, Ryan A. Koppes

**Author notes:** Correspondence and requests should be addressed to R.A.K. or N.A.

## Abstract

Excitation-contraction (EC) coupling in the heart has, until recently, been solely accredited to cardiomyocytes. The inherent complexities of the heart make it difficult to examine nonmuscle contributions to contraction *in vivo*, and conventional *in vitro* models fail to capture multiple features and cellular heterogeneity of the myocardium. Here, we report on the development of a 3D cardiac μTissue towards recapitulating the architecture and composition of native myocardium *in vitro*. Cells are encapsulated within micropatterned gelatin-based hydrogels formed via visible light photocrosslinking. This system enables spatial control of cardiac microarchitecture, perturbation of the cellular composition, and functional measures of EC coupling via video microscopy and a custom algorithm to quantify beat frequency and degree of coordination. To demonstrate the robustness of these tools and evaluate the impact of altered cell population densities on cardiac μTissues, contractility and cell morphology were assessed with the inclusion of exogenous non-myelinating Schwann cells (SCs). Results demonstrate that the addition of exogenous SCs alter cardiomyocyte EC, profoundly inhibiting the response to electrical pacing. Computational modeling of connexin-mediated coupling suggests that SCs impact cardiomyocyte resting potential and rectification following depolarization. Cardiac μTissues hold potential for examining the role of cellular heterogeneity in heart health, pathologies, and cellular therapies.

---

Ischemic heart disease remains a leading cause of death worldwide. While pharmacological interventions have improved life expectancy by mitigating key risk factors, therapeutic strategies for repairing the damaged myocardium have yet to become the clinical standard^1^. Following infarct, delivery of autologous cells, including mesenchymal stem cells (MSCs), cardiac stem cells, and endothelial progenitor cells, have yielded promising results at the benchtop, but inconsistent benefits in clinical trials^2^. To improve the efficacy of potential therapeutic strategies, new experimental models at the benchtop that enhance our fundamental understanding of the contribution of cardiac support cells are required.

In a healthy heart, cardiomyocytes (CMs) are the most abundant cells by volume, but only make up 25-35% of the total myocardial cells^3,4^. While the percentage of CMs in the heart is relatively accepted, there is a lack of consensus regarding the interstitial cell composition in the heart^4^. Current perspectives on the cardiac composition suggest the presence of fibroblasts^3–7^, endothelial cells^3–7^, smooth muscle cells^5–7^, pericytes^5,7^, Schwann cells (SCs)^5,7^, macrophages^4,5,7^, telocytes^5,7^, stem cells^5,7^, conduction cells (pacemaker cells and Purkinje fibres)^6,8^, neurons^9^, and atrial and ventricular CMs^6^. Not only is the composition diverse, but an accurate understanding of the cellular composition is critical to fully comprehend cardiac pathogenesis and aid the development of new therapeutic strategies^4,10^. Further, the role of electric-coupling of resident non-myocyte cells, such as the fibroblasts^11^, macrophages^12^, telocytes^13^, and stem cells^14^ has only recently been observed and modeled. However, due to current experimental limitations, the investigation of these cell types and their interactions require a combination of *in vivo, in vitro, and in silico* examination.

The development of robust and biomimetic *in vitro* models that mimic the architecture and heterogeneity of *in vivo* counterparts is essential for the study of fundamental biological processes. Specifically, topographical cues to promote cellular alignment has been demonstrated to promote a more biomimetic cardiac phenotype^15^. Nevertheless, while traditional two-dimensional (2D) cell culture techniques have been used for decades and contributed greatly to our fundamental understanding of cardiomyocyte function, their simplicity is often outweighed by their limitations such as atypical size and morphology, de-differentiation, and limited cell-cell contacts^16^. The development of three-dimensional (3D) cell culture has helped to overcome some of these limitations by providing more physiologically relevant environments^17^. In typical 3D culture, cells are encapsulated within extracellular matrix (ECM) like materials, most often hydrogels, to closer mimic *in vivo* microenvironment^18^. However, current 3D cell culture models are often limited to simple geometries or require the use of complex and expensive microfabrication techniques or microfluidics to form more biomimetic 3D constructs. For example, cell printing has enabled the production of complex 3D architectures *in vitro*^19^, however, the shear forces that cell experience during processing have greatly precluded the use of sensitive cell types like CMs.

The development of photocrosslinkable hydrogels facilitates an alternative to 3D printing by patterning complex shapes and structures to recapitulate *in vivo* architecture of tissues including the heart^20,21^. However, the reliance on ultraviolet (UV) light in these systems may lead to reduced cell viability^22^. Further, while photocrosslinkable materials are relatively inexpensive to synthesize, the specialized light sources required to polymerize the materials are prohibitively expensive for widespread use. More specifically, there are no *in vitro* models to investigate the impact of resident non-CM cells on cardiomyocyte function. Therefore, there remains a need to develop an alternative inexpensive approach for designing an *in vitro* 3D culture model that can be scaled to systematically examine the multicellular nature of the cardiovascular system.

Here, we describe the development of a biomimetic, 3D *in vitro* model (or cardiac *μ*Tissue) to examine the role of different cell types in the heart. Specially, primary cardiac cells were encapsulated at different ratios and variable geometries to assess the importance and function of SCs and the necessity for local alignment in the heart. To mimic the architecture and cellular alignment within the native heart a photolithography technique was used to encapsulate CMs within micropatterned gelatin methacrylate (GelMA) hydrogels via an inexpensive visible light LED (405 nm) system. Cardiac *μ*Tissue functional output was evaluated on a cell-by-cell basis using a novel algorithm to measure beats per minute as well as the degree of coordination over the tissue construct using video microscopy. In addition, functional differences between visible and UV light photocrosslinking systems for cardiac cell encapsulation were compared in patterned and unpatterned systems. As a proof of concept for the potential of *in vitro* photo-patterned cardiac *μ*Tissues to study cardiovascular health and repair, the role of exogenous SC incorporation, an abundant cell type whose function in the heart remains unknown, was experimentally examined (**Fig. 1A**). Lastly, a computational model of SC-CM coupling was developed in order to provide deeper insight on our experimental observations and the underlying SC impact on CM electrophysiology.

**Figure 1.**
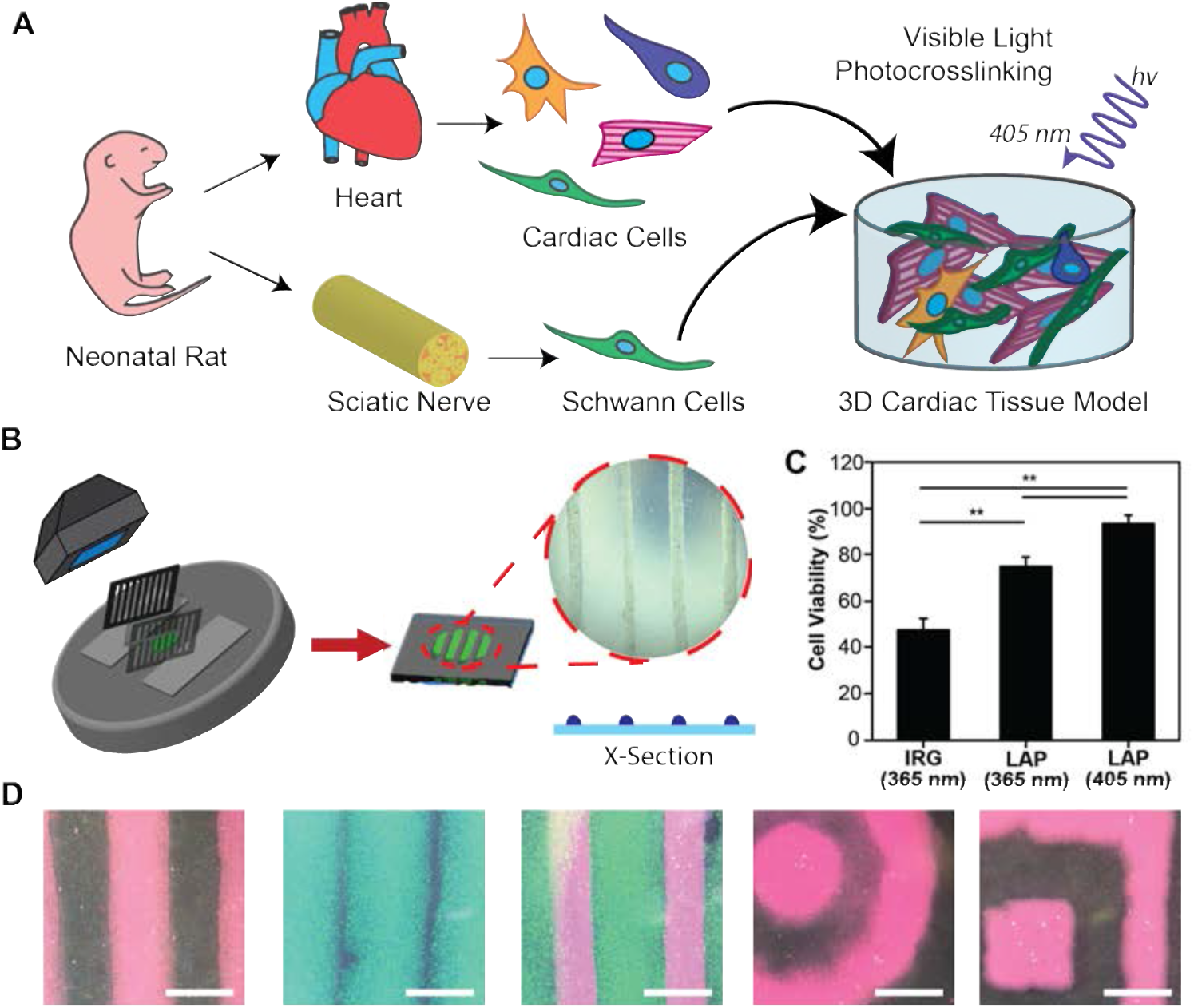
Cardiac μTissue development. **(A)** Schematic representation of the *in vitro* co-culture system to investigate the heterogeneity of the myocardium. **(B)** Schematic of photopatterning hydrogels using visible light. **(C)** Quantification of live/dead images show >90% cardiac cell viability in GelMA crosslinked using LAP with visible light (** p < 0.05). **(D)** Representative images demonstrating an ability to photopattern a range of geometries (scale = 1000 μm).

## Results

### Cardiac Tissue Models

Here, 3D cardiac tissue models (μTissues) were developed to rapidly and inexpensively synthesize cell-laden tissue constructs using commercially available materials. GelMA hydrogels were crosslinked with either a UV or visible light photoinitiator system: Irgacure using an Omnicure S2000 (365 nm) or LAP using a custom 405nm LED system (Supplemental Fig. 2A). The compressive modulus of the visible crosslinked hydrogels, measured via unconstrained cyclic compression testing, increased significantly from 1.16 ± 0.04 kPa for 5% (w/v) GelMA to 22.56 ± 1.91 kPa for 10% GelMA to 80.49 ± 2.75 kPa for 15% GelMA (Supplemental Fig. 2B). Based on these data, a 7.5% (w/v) GelMA hydrogel was selected for cell encapsulation to best mimic the stiffness of the native cardiac tissue (mean 6.8 kPa)^31^.

Primary cardiac cells were then encapsulated in patterned (500 μm wide lines with 2000 μm separation) GelMA hydrogels by controlling the areas of the precursor hydrogel that were exposed to light using laser cut black cardstock paper (**Fig. 1B**). The resulting cell-laden hydrogels exhibited significantly increased cell viability (93.44 ± 3.75%) compared to constructs formed with UV light irrespective of the photoinitiator (**Fig. 1C, Supplemental Fig. 3**). Lastly, we used photolithographic masks to fabricate a range of 3D geometries otherwise only achievable using microfluidics or 3D bioprinting (**Fig. 1D**).

### Functional Characterization of Cardiac Output

Traditionally, CM contraction *in vitro* is best quantified by electrophysiological recordings via a MEA^32^. However, measurements with planar MEAs of membrane potentials within 3D culture environments is difficult, resulting in lost recording resolution and signal-to-noise ratios. To overcome this challenge as well as mitigate the financial hurdle of an electrophysiology rig, previous groups have used video microscopy to measure cardiac frequency^32–34^, but have not considered the importance of the degree of coordination between CM contractions as an indicator for myocardium maturity. To garner a clearer sense of isolated vs. conducted contraction in our 3D *μ*Tissues, we developed an automated MATLAB algorithm to measure both CM beating frequency and degree of coordination from video recordings. Individual cells were identified within 3D hydrogels by this algorithm (**Fig. 2A**), and classified as beating CMs or a non-beating cardiac cells based upon inclusion criteria (i.e., signal-to-noise ratio, peak-to-peak frequency) described above (**Fig. 2B**). The summation of contractions was used to calculate the global average BPM for each experimental condition.

**Figure 2.**
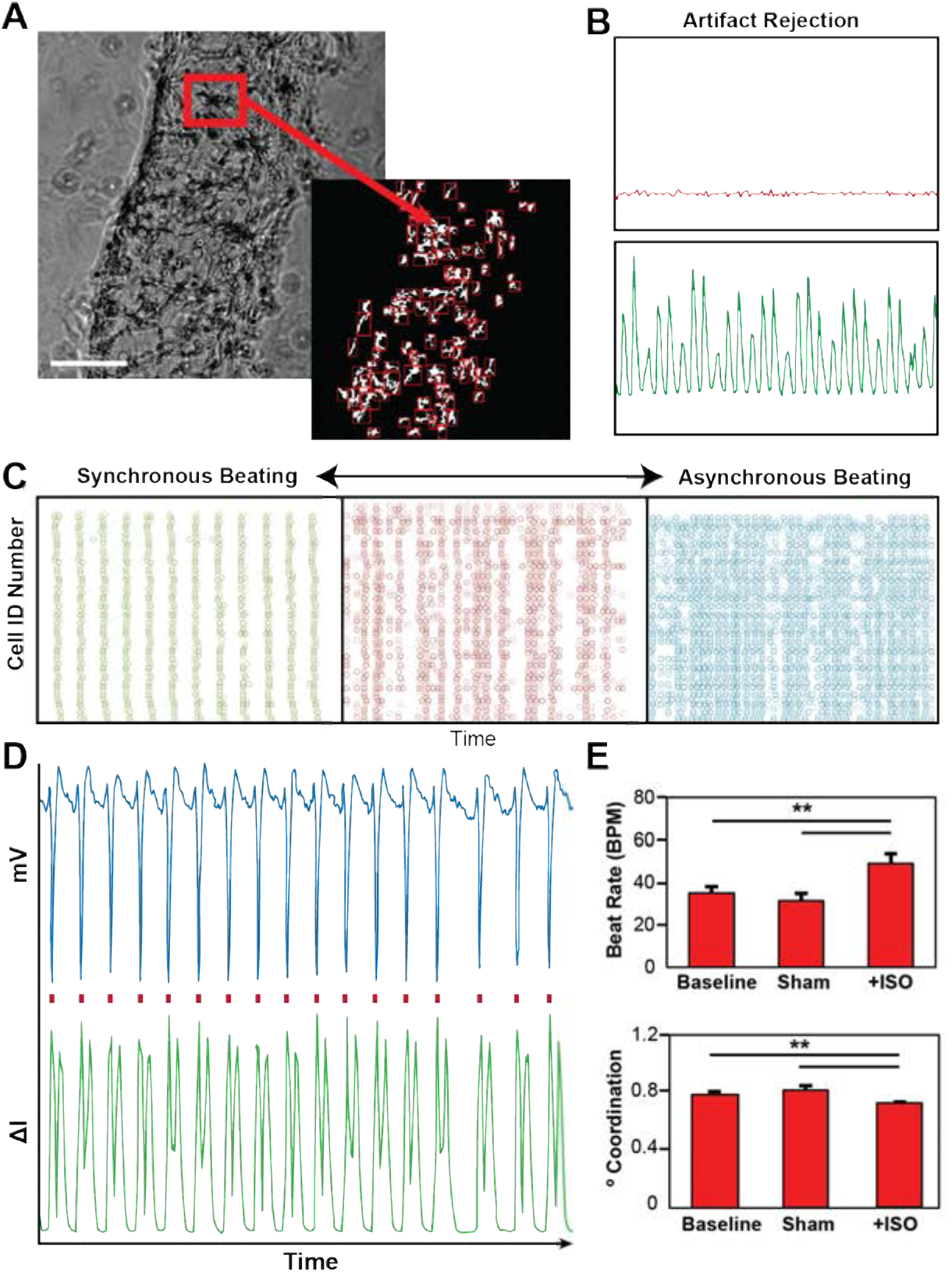
Cardiac beating quantification via video microcopy. **(A)** Representative bright field image with automated identification of encapsulated cells (scale = 200 μm). **(B)** Representative plots showing the change pixel intensity over time for both a beating and non-beating cell. **(C)** Representative spike train analog used to quantify the degree of coordinated contractions (green = high level of synchrony, red = some semblance synchrony, blue = no apparent synchrony). **(D)** Validation of beating quantification shows a one-for-one correlation with electrophysiology recordings. **(E)** Isoproterenol treatment of encapsulated cardiac cells shows an expected increase in beat rate and a decrease in synchrony measured via video microcopy (** p < 0.05).

The degree of coordination was quantified by assigning each beating CM with a unique cell ID# and plotting the timestamp of each contraction (**Fig. 2C**). To quantify coordination, each of these plots was analyzed in a similar means to the SPIKE algorithm developed for assessing dependence of action potentials in neuron populations^26^. As shown in **Fig. 2D**, this image quantification of individual cell beating has a one-for-one correlation with recordings processed using an MEA, thereby demonstrating the robustness of our custom algorithm for detecting contractions of individual cells and their interdependence in our 3D *μ*Tissues. As further validation, encapsulated cardiac cells were treated with isoproterenol, a beta-adrenergic agonist to demonstrate the utility of this algorithm to detect a changes in physiological responses^35^. A temporal increase in beating frequency (~50%) and decrease in coordination (~10%) was detectable with this video quantification tool (**Fig. 2E**).

### Impact of Patterning and Photoinitiator on 3D *μ*Tissues

Spontaneous beating was recorded over a period of nine days for cardiac *μ*Tissue constructs formed using both visible and UV light and for patterned and unpatterned hydrogel samples. Similar to previous findings^33^, no beating was observed when only enriched populations of CMs were encapsulated within the 3D *μ*Tissues (data not shown). *μ*Tissues crosslinked with visible light (LAP), on average, exhibited on average a 25% increase in rates of beating for all measured time points compared to Irgacure^®^ controls (**Fig. 3A-C**). Interestingly, there were no differences in the degree of coordination were apparent when comparing crosslinking systems irrespective of patterning (**Fig. 3B-C**). However, regardless of the crosslinking system implored, the inclusion of ACCs within patterned hydrogels led to spontaneous beating on day 3 post encapsulation vs. day 4 post encapsulation for unpatterned samples (**Fig. 3A, Fig. 3D**). Lastly, patterned 3D *μ*Tissues exhibited a statistically significant (p < 0.0001) increase in degree of coordination compared to unpatterned hydrogels throughout the entire study (**Fig. 3E-F**).

**Figure 3.**
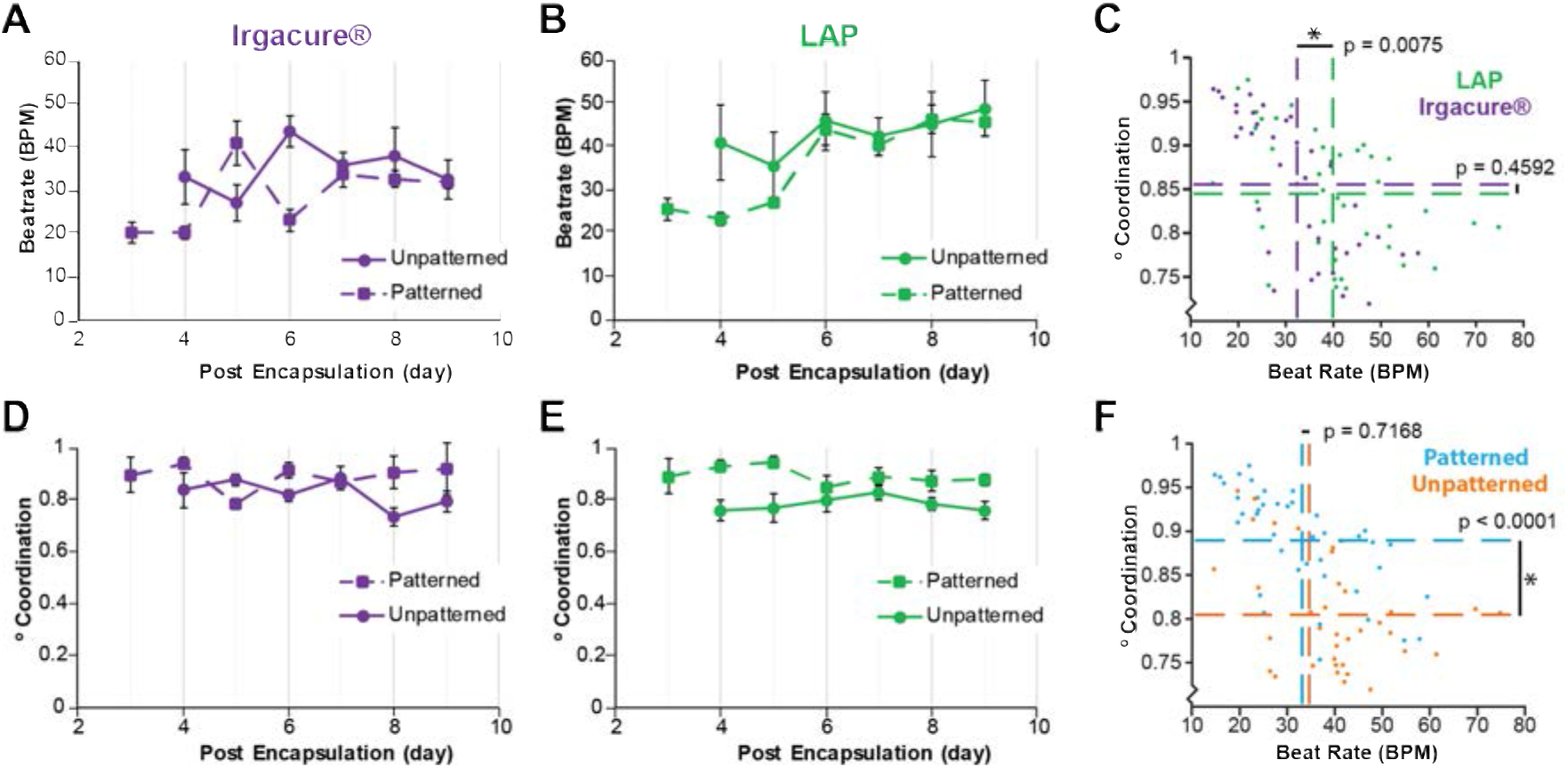
Model development informed by cardiac output quantification. **(A)** Quantification of beat rate for cardiac cells encapsulated in unpatterned and patterned hydrogels over time using a UV (Irgacure^®^ and 365 nm) or **(B)** a visible (LAP and 405 nm) or crosslinking system. **(C)** Multi-way ANOVA showing statistical differences between the visible and UV crosslinking systems for beat rate, but not coordination. **(D)** Quantification of beating coordination for cardiac cells encapsulated in patterned or unpatterned hydrogels formed using a UV (Irgacure^®^ and 365nm) or **(E)** visible light (LAP and 405nm) crosslinking system. **(F)** Multi-way ANOVA showing statistical differences between patterned and unpatterned samples for beating coordination, but not beat rate.

### Impact of SC incorporation in the 3D *μ*Tissues

To validate the use of cardiac *μ*Tissues for further understanding the role of non-myocyte support cells in the heart, functional outputs and morphological differences were assessed with the inclusion of exogenous SCs, which endogenously are the 3^rd^ most abundant interstitial cell in the heart^5^. Enriched CM and ACC populations were characterized via immunocytochemistry to quantify the endogenous cell composition as a baseline of isolation heterogeneity. As expected, an increased ratio of CMs, identified by sarcomeric *α*-actinin, were present in the enriched CM suspension (Fig. 4A-C). The remaining cells in the both the enriched CM and ACC cultures were identified by staining for fibroblasts (CD90), smooth muscle cells (α-smooth muscle actin), and SCs (S100). These cell types were investigated due to pervious reports suggesting their abundance in the myocardium^3,4^. However, while, immunostaining demonstrated that both the enriched CM and ACC cultures contained heterogeneous cell populations, this analysis only provides a glimpse of all the cells found in the heart. The proportion of viable S100 positive cells isolated from the heart was 19.61 ± 4.08%, which is approximately double of the composition previously reported for the human atria^5^.

**Figure 4.**
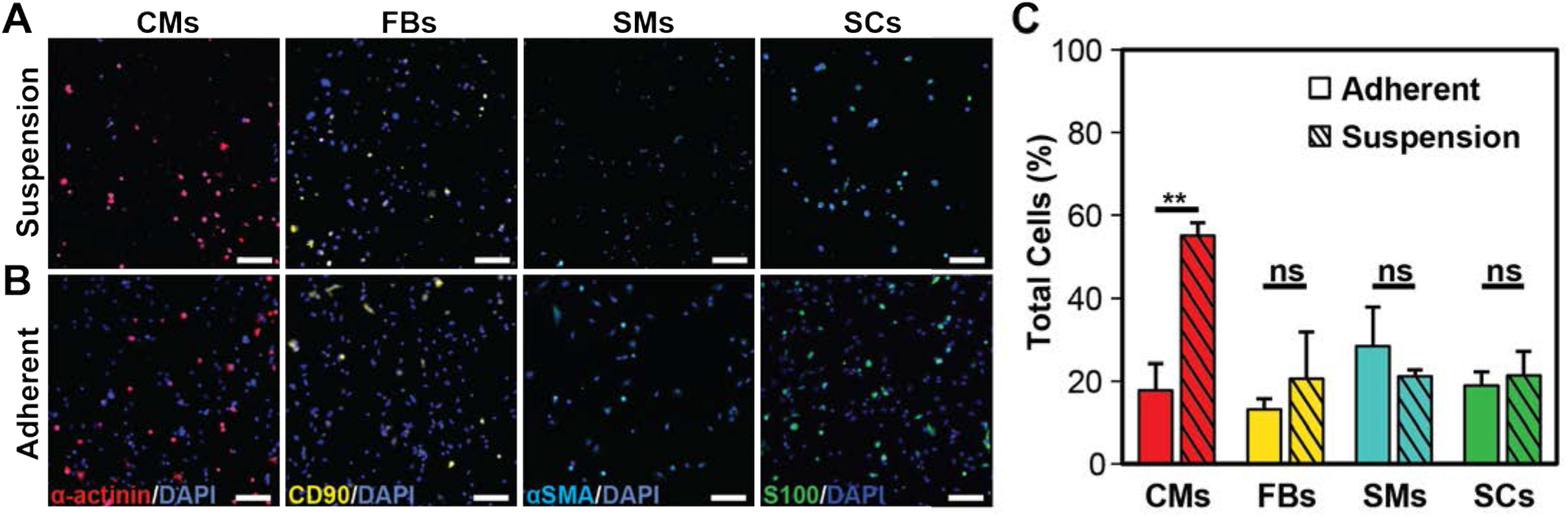
Immunofluorescent quantification of cardiac cellular composition. **(A)** Representative suspension and **(B)** adherent cultures grown on fibronectin coated cover slides for 12-18h post tissue dissociation (scale = 100 μm). Cells were stained for markers for cardiomyocytes, sarcomeric alpha-actinin (red), fibroblast, CD90 (yellow), smooth muscle cells, alpha smooth muscle actin (cyan), Schwann cells, S100 (green) and cell nuclei, DAPI (blue). **(C)** Quantification of immunofluorescent images show that the suspension culture contains an enriched population of cardiomyocytes, with no statistical differences been other cell populations examined. (** p < 0.05, ns: not significant p > 0.05)

Injected SCs have demonstrated therapeutic effect on cardiac function following infarction in rodent models^36,37^. To more closely examine how SCs effect CMs, exogenous SCs were encapsulated along with endogenous cardiac cells in our 3D μTissues. Here, SCs from the sciatic nerve were used to ensure a high concentration and avoid a potential loss of phenotype that has previously been reported for cardiac support cells during expansion^16^. Inclusion of SCs within the cardiac *μ*Tissues led to a significant 2-fold increase (p < 0.0001) in the beating frequency of CMs as compared to controls at all measured time points (**Fig. 5A**). Additionally, the inclusion of SCs decreased the degree of coordination in the 3D *μ*Tissues from average of 85.10 ± 0.98% to 76.76 ± 0.99% (**Fig. 5B**). These trends were also observed for cardiac cells with/without SCs when cultured on a 2D scaffold (**Supplemental Fig. 4**). There were no observable morphological differences of CMs within the 3D hydrogel cultures after the incorporation of SCs (**Fig. 5C-D**), yet in 2D cultures, CMs co-cultured with SCs possessed an aspect ratio of 4.1 ± 1.8 compared to 1.6 ± 0.5 for controls without SCs (**Fig. 5E-G**).

**Figure 5.**
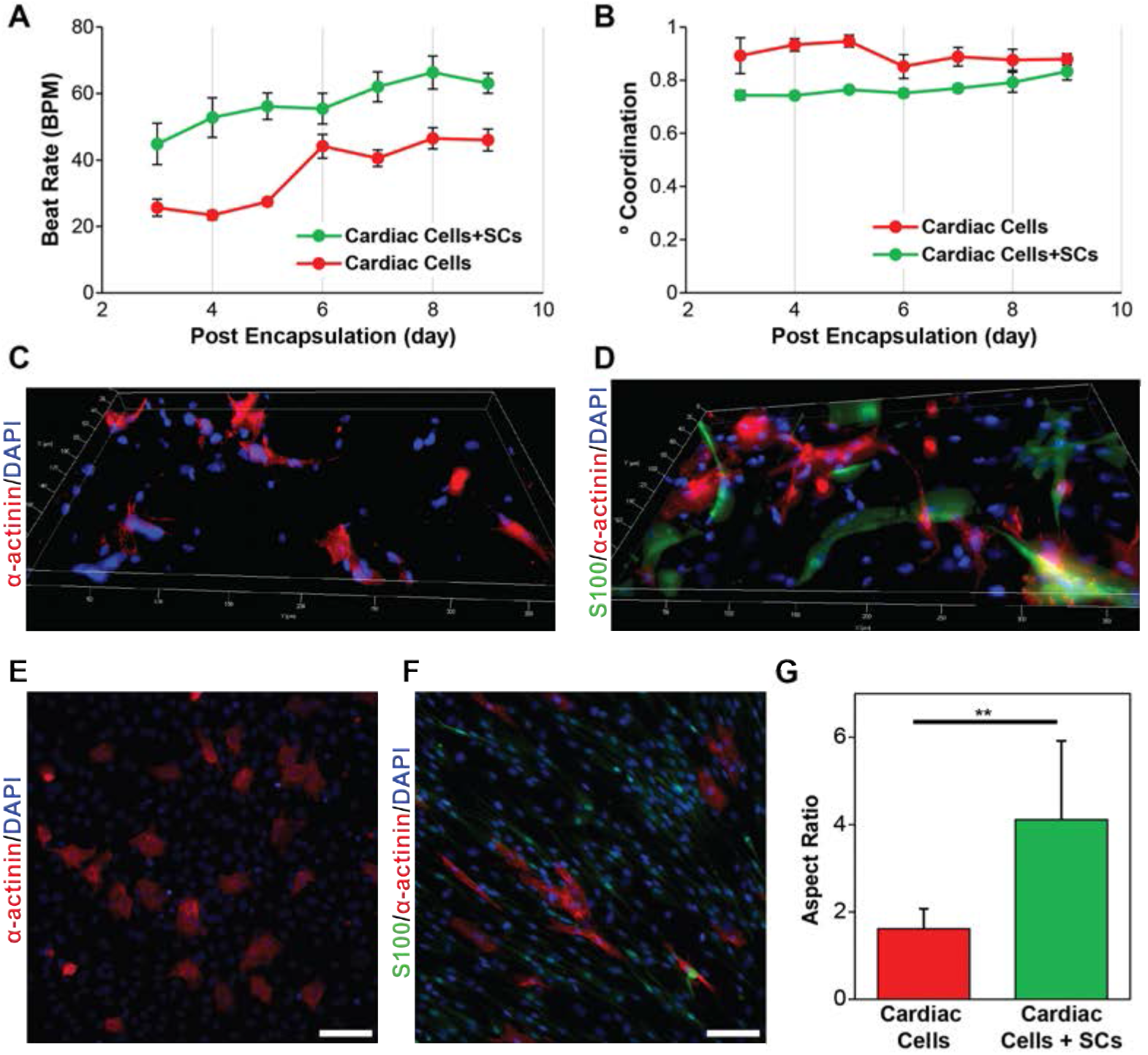
Role of SCs in the in 2D and 3D cardiac cell cultures. **(A)** Quantification of beat rate and **(B)** degree of coordination for 3D μTissues over time showing that the inclusions of SCs leads to an increase and beat rate and decrease if coordinated contracts. **(C)** Representative immunofluorescent images of cardiac cells culture without and **(D)** with the inclusion of exogenous SCs encapsulated in 3D μTissues. **(E)** Representative immunofluorescent images of cardiac cells culture without and **(F)** with the inclusion of exogenous SCs in 2D (scale = 100 μm) and **(G)** their respective cell aspect ratio quantification (** p < 0.05).

### Electrical Pacing of 3D *μ*Tissues

Electrical stimulation (ES) to pace the 3D *μ*Tissues provided better understand how SCs may influence contraction of the myocardium. ES was applied to the *μ*Tissues on day 5 post encapsulation via carbon rod electrodes as previously reported^27^. Rates of beating were measured for video recording taken at different frequencies and voltages. Prior to ES, encapsulated cardiac cells had an average BPM of 45.27 ± 2.16. Ramping the voltage from 1V to 3V to 5V led to an increased response that trended toward the applied frequency for 2 (117.19 ± 8.85) and 3 Hz (153.11 ± 6.95) at 5V (**Fig. 6A**). The cardiac *μ*Tissue constructs were unable to be paced at 0.5 or 1 Hz for any of the tested voltages. As the rate of spontaneous beating was close to the 1 Hz pacing, the effects of electrical stimulation in this range were indeterminable. Whereas, the incorporation of SCs into these constructs (average BPM of 60.50 ± 9.51) completely prevented electrical pacing at any of the tested frequencies and voltages (**Fig. 6B**). Immunocytochemistry of the co-culture suggest that SCs ability to insulate CMs may in part be due to the formation of connexin junctions between SCs and CMs (Fig. 6C). Lastly, with the addition of a connexin blocker, heptanol, only a small percentage of CMs could be electrically paced (**Fig. 6D**), thereby further suggesting CM-SC coupling.

**Figure 6.**
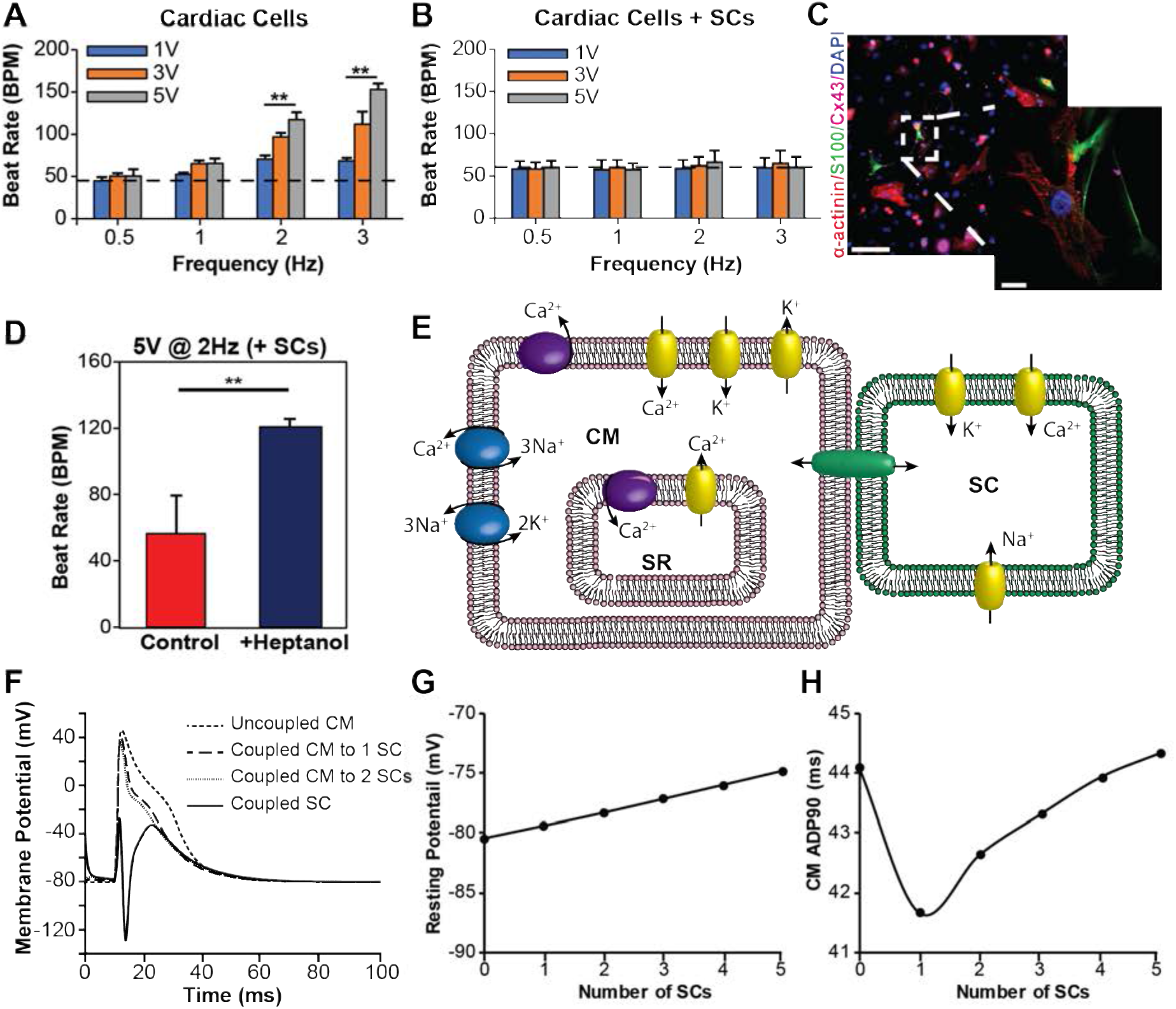
External electrical pacing of 3D μTissues and computational modeling on CM-SC coupling. **(A)** Cardiac response to electrical stimuli at 0.5, 1, 2, and 3 Hz with increasing voltages from 1V to 3V to 5V shows cardiac *μ*Tissues can be paced (** p < 0.05), but **(B)** the inclusion of SCs prevents this electrical pacing (dashed line shows the average beat rate prior to electrical stimulation). **(C)** Representative immunofluorescent image showing the presence of connexin43 junctions (purple) between CMs (red) and SCs (green) (scale = 100 μm, inlay scale = 10 μm). **(D)** Electrical pacing at 2Hz and 5V of cardiac cells with exogenous SCs following heptanol treatment to block CM-SC coupling (** p < 0.05). **(E)** Schematic of a Hodgkin-Huxley model for CM-SC coupling. **(F)** Computational model of CM depolarization for one CM coupled to increasing numbers of SCs overplayed with a coupled SC membrane potential. **(G)** Changes in CM membrane potential and **(H)** APD90 with increased SC coupling.

### Modeling of CM-SC Coupling

As evidence by our visual analysis of 3D *μ*Tissues contractility, SCs influenced the electrophysiological properties of CMs *in vitro*. To further explore this interaction, we modeled CM-SC coupling *in silico* (**Fig. 6E**). We first developed a simple SC model by fitting published SC voltage clamp data for a type I potassium channel^28^, a L-type calcium channel^29^, and a sodium channel^30^. Equations for maximum current, channel conductance, activation and inactivate coefficients, and time constants vs. membrane potential were used to simulate voltage clamp experiments for each SC channel (**Supplemental Fig. 5–7**). Coupling the simulated SC with a model CM^11^ suggested that SCs increase the CM resting potential and accelerates repolarization (**Fig. 6F-H**). *In vivo*, a decreased APD90 increases the frequency at which a CM can be depolarized^12^, thereby increasing the maximum beating frequency. However, contrary to our results from electrical pacing, a higher resting potential did not reduce the stimulation threshold necessary to facilitate depolarization.

## Discussion

Our developed biomimetic *in vitro* 3D *μ*Tissues hold potential to measure functional outputs as cardiac cellular composition is manipulated. This system approaches the resolution of cell printing without subjecting sensitive primary CMs to the temperatures and shear stresses often associated with bioprinting conditions^38^. CMs were encapsulated in a patterned ECM-like material, GelMA, to mimic the native cardiac chemical and architectural cues^31,34^. GelMA allows tunability to mimic native mechanics, possesses protein composition to native myocardium ECM as well as a high availability of cell binding domains, and can be processed into 3D geometries^20^. However, traditional GelMA crosslinking requires high intensity UV and cytotoxic photoinitiators^20^. Thus, alternative visible light crosslinking systems have been recently developed^20,22,39,40^.

EosinY/TEA is a type II photoinitiator that has been demonstrated to support high cell viability for encapsulated 3T3 fibroblasts^22^, but poor viability for primary CMs^24^ in crosslinked GelMA hydrogels. Type II photoinitiators also require longer photocrosslinking exposure times to achieve the same mechanical properties as type I photoinitiators^40^, which may lead to decreased cell viability due to the increase in free radicals^41^. Here, we investigated the use of LAP, a commercially available visible light type I photoinitiation, for the development of our cardiac *μ*Tissues. As expected, GelMA hydrogels formed with either photoinitiator (LAP and Irgacure^®^) exhibited similar mechanical properties (**Supplemental Fig. 2B**), within the ranges previously reported^20^. Primary cardiac cells encapsulated in GelMA hydrogels using LAP as a photoinitiator and visible light demonstrated increased cell viability as compared to those encapsulated in hydrogels formed by using UV light and/or Igacure^®^, as reported previously in other cell studies^40^.

Correlating in-the-dish measures of contractility to cardiac output and health is paramount for the long-term utility of *in vitro* cardiac models. Previously, beating frequency, degree of coordination, and conduction velocity have been exclusively measured electrically^15^. However, methods for intra and extracellular recordings require expensive equipment and highly specialized expertise. Furthermore, use of these techniques with 3D *μ*Tissues provides additional technical challenges. Attempts to overcome the limitations of using electrophysiology-based approaches have yielded some success by measuring contractions via video microscopy^32–34^ or cantilevers^42–44^. In each of these cases, beating frequency is measured by counting the global changes in intensity, size, and/or force measurements over a set duration. However, these techniques fail to directly measure variability across individual beating cells, and therefore the quantification of coordination and conduction velocity remain out of reach for these techniques. Here, we developed a custom processing technique that allows for cardiac function to be assessed on a per cell basis that renders comparable results to traditional electrophysiological recording devices (**Fig. 2**). Our algorithm is capable of calculating conduction velocity, however, accurate measurements rely on frame rates faster than 200 frames per second to accurately measure differences between cells in the same field of view, which currently lies beyond the capabilities of most camera systems.

As a proof of concept for examining the efficacy of our 3D *μ*Tissues for the investigation of the complex cellular composition of the heart, we sought to understand how SCs might affect cardiomyocyte phenotype. SCs are nonneural support cells that play an essential role in neuron signaling and regeneration^45^. SCs have been evaluated at the bench as a cell therapy for cardiac failure^37,46^ and make up a substantial composition of the heart^5,7^. Often cells isolated from the heart are characterized as either CMs or cardiac FBs^22,24,33,34,47,48^, but in fact these populations are heterogeneous and contain fewer than 20% FBs. For this reason, we aimed to establish a new nomenclature for this cell isolation protocol in which suspension cells are referred to as enriched CMs and the adherent cells are referred to as adherent cardiac cells (ACCs). Further, the presence of these ACCs is critical for developing functional engineered cardiac tissues from iPSCs^10^.

Exogenous SCs were incorporated into our 3D *μ*Tissues to elucidate the impact of SCs in cardiomyocyte phenotype and function. CMs co-cultured with SCs in standard 2D conditions exhibited an increased aspect ratio (between 5 and 9.5; **Fig. 5A-C**), typical of a more mature CM phenotype^49^. Qualitatively, CMs cultured with SCs present with more clearly defined z-disks, evidenced by localization of -actinin, elongation of nuclei, and a possible an increase in bi-nucleation (**Supplemental Fig. 8**): indicators of CM differentiation and maturation^50,51^. In response to peripheral nerve damage, non-myelinating SCs provide topographical cues that influence the rate and direction of neurite outgrowth^52^. Monolayers of SCs also exhibit local alignment that impacts the growth of dissociated sensory neurons *in vitro*^53^, suggesting that CMs may exhibit a higher aspect ratio by a similar mechanism. Peculiar to this observation of increased CM differentiation, the presence of exogenous SCs increased the average beats per minute as well as a decrease the degree of coordination in engineered cardiac *μ*Tissues (**Fig. 5F-G**), similar to higher BPM observed during development^54^. However, in addition to providing guidance cues, SCs also overexpress matrix metalloproteinases (MMPs) following a nerve injury to help clear cellular debris^55,56^. Therefore, due to an abundance of MMP-sensitive degradation sequences in gelatin^57^, the inclusion of SCs may result in lower mechanical properties of our *μ*Tissues over time, which may affect the cardiac phenotype and resulting cardiac output^31^. This reduction in mechanical stability was confirmed observationally by the increase degradation and delamination of the *μ*Tissues from the glass slides after 9 days (**Supplemental Fig. 9**). Nevertheless, similar cardiac output trends were observed when cells were cultured on 2D scaffolds comparted to those cultured in 3D, thus ruling out any confounding effects of material degradation (**Supplemental Fig. 4**).

Developing mammalian hearts beat at higher rates due a higher ratio of connective tissue to muscle or ACCs to CMs, immature T-tubule and sarcoplasmic reticulum, and a higher concentration of plasma Calcium^58^. CM contraction is predominantly regulated by intracellular Ca^2+^ concentration cycling and a resultant membrane depolarization, that in mature CM constructs originates with an atrial pacemaker and progresses to surrounding cells with a high degree of coordination. Non myelinating SCs exhibit excitable properties including regulating Ca^2+^ in neuromuscular junctions^59^ and K^+^ in the axonal microenvironment^60^. While SC presence in the heart has just recently been highlighted, their role on plasma and cytosol ion dynamics in the myocardium remains unknown.

Interestingly, when external electrical stimulation was applied to our 3D *μ*Tissues, SCs prevented the CMs from being paced. We hypothesize that SCs, which are known to express connexin proteins^61^, form connexin junctions with some CMs (**Fig. 6C**), therefore increasing the resting potential and allowing for a more rapid depolarization and subsequent contraction and expansion. This mechanism of CMs forming connexin junctions with other cells is demonstrated by a recent report in which cardiac macrophages are shown to increase the CM resting potential^12^. While SCs maintain a higher resting membrane potential compared to CMs (−40 mV for SCs^28,30^ vs. −80 mV for CMs^11^), SCs also contain ion channels for calcium and sodium in addition to similar potassium channels to those in cardiac macrophages. Therefore, it is expected that in addition to an increased CM resting potential with increased SC coupling, the CM electrophysiological properties will be significantly altered. Executing the CM-SC *in silico* model revealed that the initial depolarization of the SC membrane with CM depolarization led to an overcorrection to repolarize the SC membrane, which helped to quicken the CM repolarization. However, with increased coupling, the rate at which a CM is repolarized returned to its initial APD90, thereby suggesting an enhanced ability for CMs coupled with SCs to better regulate their membrane potential, while initial resting potential continues to increase. Lastly, when treated with a general connexin blocker, heptanol^62^, some CMs exhibited correspondence to external, pulsatile electrical stimulus. This finding suggests that connexins are involved in SCs ability to electrically couple to CMs, similar to that observed with fibroblasts^11^ and macrophages^12^. However, the mechanism which leads to increased beating and diminished coordination requires further investigation.

By studying tissue engineering or cell therapies *in vitro* we aim to identify the underlying mechanisms of healthy and diseased cardiac tissue with a goal of improving current therapeutic strategies. Specifically, our 3D *μ*Tissues provide a biomimetic microenviroment that recapitulates the *in vivo* structure, architecture, and cellular composition of the heart, which is critical for the development of cell and tissue engineering therapeutic strategies^63^. Furthermore, this platform can be exploited to better model human physiology and pathology through the incorporation of stem cell derived cardiac cell populations. These approaches in conjunction with computational models, quantitative cardiac outputs, and low capital expensive will hopefully lead to microphysiological systems, which can be utilized to increase translation of novel therapies to the clinic. All algorithms for both analysis and modeling can downloaded from http://www.northeastern.edu/lnnr.

## Methods

### GelMA synthesis and hydrogel fabrication

GelMA, derived from fish gelatin, was synthesized as previously reported^23^. In brief, 8% (v/v) methacrylic anhydride (Sigma) was added to a 10% (w/v) fish gelatin (Sigma) solution in Dulbecco’s Phosphate Buffered Saline (DPBS, Sigma). The product of this reaction was dialyzed (Spectra/Por 12-14kD, Fisher Scientific) using distilled water for 1 week, and then lyophilized for use on demand. GelMA hydrogel precursor solutions (7.5% (w/v)) were prepared in complete culture medium [Dulbecco’s Modified Eagle Medium (DMEM) without phenol red (Sigma) supplemented with 10% fetal bovine serum (Sigma), 2 mM L-glutamine (Gibco), and 50 units/mL penicillin/streptomycin (Gibco)] with 0.5% (w/v) 2-hydroxy-1-(4-(hydroxyethoxy) phenyl)-2-methyl-1-propanone (Irgacure^®^ 2959, CIBA Chemicals) or 0.5% lithium phenyl-2,4,6-trimethylbenzoylphosphinate (LAP, Biobots) as photoinitiators. Hydrogel precursors were then photocrosslinked by exposure to UV light (365 nm) with an Omnicare S2000 (Excelitas Technologies) for Irgacure^®^ containing precursor hydrogels or visible light (405 nm) with a 10W LED (QUANS) for LAP containing precursor hydrogels (0.25 seconds of 10 mW/cm^2^ light exposure per *μ*m of hydrogel thickness).

### Mechanical characterization

The compressive modulus of each hydrogel formulation (5%, 10%, and 15% (w/v) polymer concentration for UV and visible light crosslinked hydrogels) was examined using an ElectroForce mechanical tester (TA instruments) with a 1000 gr load cell. Hydrogel samples were prepared in custom polydimethylsiloxane (Sylgard) molds (cylinders of 6 mm diameter; 4 mm height). Hydrogels were loaded between two compression platens, and cyclic uniaxial compression tests were conducted at 0.5 Hz (10 cycles). Compression displacement and load for each cycle were recorded using WinTest7 software. The compressive modulus was calculated as the tangent slope of the linear region of the stress-strain curves between 0.1 − 0.25 stain level. Three independently prepared samples for each formulation were measured to quantify the compressive modulus.

### Primary cardiac cell isolation

Primary CMs and adherent cardiac cells (ACCs) were isolated from two-day old (p2) Sprague-Dawley neonatal rat (Charles River) hearts, following a modified protocol by Noshadi *et el*.^24^ and approved by Northeastern’s Institutional Animal Care and Use Committee (IACUC). In brief, the thorax was opened and the heart was removed. The major veins and atria were then removed so that only the left and right ventricles remained. The ventricles were cut into 3-4 pieces and stored in 0.05% (v/v) trypsin (Gibco) in Hank’s balanced salt solution (HBSS, Gibco) at 4 °C with continuous shaking overnight. The following day, the ventricle pieces were removed from the trypsin and subject to sequential collagenase treatments (305 units Collagenase II (Gibco) in HBSS) at 37 °C to dissociate the connective tissue and collect cardiac cells. Cells were then filtered through a 70-*μ*m cell strainer (Falcon) and pre-plated in a tissue culture flask (T-175, Corning) with complete culture medium to enrich the CM population by differential adhesion. After one hour in standard cell culture conditions (37 °C, 5% CO_2_), any unattached cells were considered to be CMs, while those attached cells were deemed ACCs. Each cell population was then counted and used for experimentation within one hour following completion of the pre-plating enrichment.

### SC isolation

Primary SCs were isolated from p2 Sprague-Dawley neonatal rat sciatic nerves using established protocols^25^. Sciatic nerves were harvested and kept in complete culture medium on ice for a maximum of 4 h. Dissected nerves were minced into 1-2 mm pieces under sterile conditions and incubated in a 6-well plate using complete culture medium in standard conditions. Tissue were then transferred to a new dish after visual confirmation of fibroblast migration. Three to four days after SC migration, cells were cultured with complete medium supplemented with 10^-5^ M cytosine arabinoside (ARA-C, Sigma) for 72 h to remove remaining highly mitotic fibroblasts. Next, a complement-mediated cell lysis was used to eliminate remaining fibroblasts. Cells were detached with 0.25% (v/v) trypsin/EDTA (Corning) and pelleted at 200 g for 5 min. Fibroblasts were targeted by re-suspending in 1 mL anti-CD90/Thy 1.1 (diluted 1:500 v/v in DMEM, Cedar Lane Labs) and incubated under standard conditions for 30 min. Treated cells were pelleted, resuspended in 1 mL rabbit complement, and incubated for 30 min with standard conditions to selectively lyse the fibroblasts. After incubation, cells were centrifuged, re-suspended in SC maintenance medium (complete medium supplemented with 6.6 mM forskolin (Sigma) and 10 *μ*g mL^-1^ bovine pituitary extract (Corning)), and cultured in a flask for expansion. Lysis was repeated if fibroblast impurities remained. SC purity was assessed using anti S100 (DAKO). Maintenance medium was changed every other day and SCs were passaged before 100% confluency until P10.

### 3D cell encapsulation

Equal parts of enriched CMs and ACCs (1:1), or CMs, ACCs, and SCs (1:1:1), were mixed with the hydrogel precursor solution (7.5% GelMA (w/v) and 0.5% (w/v) Irgacure^®^ or LAP) at a density of 1.5×10^7^ cells/mL. Approximately 10 *μ*L cell-laden gel precursor solution was placed between a 180 *μ*m-tall spacer and a 3-(trimethoxysilyl) propyl methacrylate (ACROS Organics) coated glass slide, followed by light exposure through a laser cut black cardstock paper photomask (500 *μ*m lines) to form patterned cell-laden hydrogels. Samples were incubated for 10 days in standard culture conditions with medium replaced every other day.

### Cardiac cell viability

Cell viability within 3D cardiac *μ*Tissues was determined via a LIVE/DEAD^®^ viability/cytotoxicity kit (Life Technologies) per instructions from the manufacturer. Live (green) and dead (red) cells then were imaged by using an inverted fluorescence microscope (Zeiss Axio Observer Z1). Cell viability was calculated by counting the number of the live cells divided by total cell number using ImageJ software. The experiment was repeated in triplicate, with a minimum of five images per sample utilized for analysis.

### Cardiac beating quantification

Individual CM contractions within the 3D *μ*Tissues were quantified with a custom MATLAB (Mathworks) code to calculate beats per minute (BPM) on a cell-by-cell basis using video microscopy. Cardiac cells were recorded at 30 frames per second with phase contrast on a Zeiss Axio Observer at 20x with an incubation chamber (37°C and 5% CO_2_). Raw video files were exported as AVIs (M-JPEG compression, 90% quality) and imported into MATLAB for analysis. Regions of interest (ROIs) were identified from the first frame of the video recording as objects between 75 μm^2^ - 1000 μm^2^ in size (**Supplemental Fig. 1A**). To quantify BPM, the sum differences in frame-to-frame pixel intensity were measured for each ROI. Inclusion criteria for a beating CM was: 1) peak amplitude was greater than one standard deviation above the mean, 2) negative spikes were less than two standard deviations below the mean, and 3) the frequency of spikes was below 6 Hz (**Supplemental Fig. 1B-C**). After passing these prerequisites, beating was recorded as the number of spikes found in inclusion criteria 1. However, for high frame rate movies (frames per second > 10), both CM contraction and relaxation may result in recorded spikes (**Supplemental Fig. 1D**). To record only a single beat per contraction, the mean inter-spike-interval was calculated for each cell and spikes occurring below this value were disregarded. The average BPM was then calculated as the mean number of contractions for all ROIs in field of view (m > 20) multiplied by 60 and divided by the video length from a minimum of three samples per condition (n > 3).

To validate of accuracy and precision of this algorithm for measuring CM beating, simultaneous video microscopy (Axio Examiner at 40x) and electrical potential via a multi-electrode array (MEA, multichannel systems) recordings were acquired and compared. Timestamps for each contraction from the identified CMs were assigned a unique identification number to gather a quantification on the degree of coordinated contraction in the *μ*Tissue models. Specifically, timestamped contraction data was imported into a modified algorithm originally developed to investigate neuronal activity^26^, and values of one minus the SPIKE-distance, an estimator of the similarity between spike trains, were reported as a means to quantify degree of coordinated CM contractions.

### External electrical pacing

Electrical pacing of CMs was assessed using a chamber designed to apply pulsatile electrical stimuli^27^ (50% duty square waves) to patterned cell-laden hydrogels at increasing frequencies (0.5, 1, 2, and 3 Hz) and voltage (1, 3, and 5 V) for 30 s. Alternating electrical stimulation was applied via two carbon rod electrodes mounted in a glass petri dish 2 cm apart. Pulsatile electrical signals were applied using a function generator (Agilent) connected to each carbon electrode with platinum wires. Alternating current (AC) stimulation was implemented to minimize hydrolysis. The chamber was filled with complete medium and each cell-laden hydrogel *μ*Tissue was aligned along the axis of the applied electric field between the two electrodes. The battery of conditions was randomized across samples to negate influence of testing order, and a minimum of five samples were examined at all frequencies and voltages.

### Immunocytochemistry

Cell-laden *μ*Tissues were fixed with 4% paraformaldehyde (30 min), permeabilized with 0.1% X-100 Triton (20 min), then blocked with 5% goat serum (>12 h) in DPBS. After overnight blocking, samples were incubated in 1:400 rabbit anti S100 (DAKO, Z0311), 1:200 mouse anti sarcomeric α-actinin (Abcam, ab9465), mouse anti CD90, and/or 1:200 goat anti connexin-43 (Abcam, ab87645) in blocking solution overnight at 4 °C. Cell-laden hydrogels were then washed thrice with DPBS and anti-rabbit, anti-mouse, and anti-goat secondary antibodies (1:200 in goat serum) were added overnight at 4 °C. Samples were rinsed with DPBS and mounted on cover slides with ProLong^®^ Gold Antifade with 4′,6-diamidino-2-phenylindole (DAPI). Lastly, the constructs were imaged using an inverted fluorescence microscope (Zeiss Axio Observer Z1).

### Computational model of SC-CM Coupling

SC electrophysiological properties were mathematically simulated by curve fitting published voltage clamp data to a Hodgkins-Huxley model using MATLAB. Specifically, type I potassium^28^, L-type calcium^29^, and the sodium currents^30^ were incorporated into this model. The electrophysiology properties of a beating CM coupled to a SC was modeled as previously described^11^. Model parameters are expanded on in **Supplementary Methods**.

### Statistical analysis

All data was first determine if it was normally distributed using the Lilliefors test in MATLAB. Normally distributed data was compared in MATLAB by using a student-t test for data sets containing only two experimental conditions (**Fig. 6D**), a one-way ANOVA for experiments with more than two conditions (**Fig. 1C, Fig. 2E, Fig. 4C, Fig. 6A-B**), and a multi-way ANOVA for comparisons across multiple days (**Fig. 3, Fig. 5A-B**). Non-normally distributed data was compared in MATLAB with the Kruskal-Wallis test (**Fig. 5G**). Error bars represent the mean ± standard deviation of measurements (**p < 0.05).

## Acknowledgements

R.A.K. and N.A. acknowledge Northeastern University and the startup fund provided by the Department of Chemical Engineering, College of Engineering at Northeastern University. R.A.K. further acknowledges the support from the National Institutes of Health (NIH, R21EB025395-01) and N.A. acknowledges the support from the American Heart Association (AHA, 16SDG31280010) and the NIH (R01-EB023052; R01HL140618).

## Author contributions

J.R.S., N.A., and R.A.K. conceived the project. J.R.S. synthesized and characterized materials, isolated neonatal rat cardiac cells, stained and imaged all samples, developed the MATLAB algorithm, and wrote the electrophysiology stimulation. J.R.S. completed the *in vitro* cell studies with assistance from J.A. D.D. isolated and purified and SCs from neonatal rats under the advisement of A.N.K. A.N.K provided intellectual input and advice. J.R.S. and R.A.K. analyzed the results, prepared the figures, and wrote the manuscript. All authors edited and provided feedback on the manuscript. R.A.K. and N.A. supervised the work.

## Competing finical interests

The authors declare no competing interests.

## Additional Information

**Supplementary Information** accompanies this paper attached below.

**Competing finical interests:** The authors declare no competing interests.

## Supplementary Information

### Methods

#### Electrophysiological Model

The CM transmembrane voltage was modeled as:

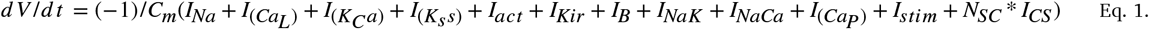

where *V* is the CM transmembrane voltage, *t* is time, *C_m_* is a CM membrane capacitance, *I_Na_* is a sodium current, *I_CaL_* is a L-type calcium current, *I_KCa_* is a calcium activated potassium current, *I_Kss_* is a steady state potassium current, *I_act_* is a hyperpolarization activation current, *I_Kir_* is an inward rectifier current, *I_B_* is a background current, *I_NaK_* is a sodium-potassium pump current, *I_NaCa_* is a sodium-calcium pump current, *I_CaP_* is a sarcolemmal Ca pump current, *I_stim_* is a stimulus current, N_SC_ is the number of SCs forming junctions with a CM, and *I_CS_* is the current from the CM-SC coupling. The *I_SC_* current was defined as:

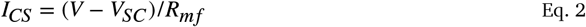

where *V_SC_* is the SC transmembrane voltage and *R_mf_* is the coupling resistance. All other currents were defined as previous described by Sachse et al^11^. The SC transmembrane voltage was modeled as:

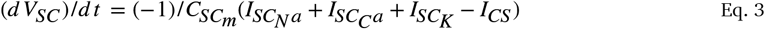

where *C_SC_m_* is a SC membrane capacitance, *I*_*SC_N*a_ is a SC sodium current, *I_SC_Ca_* is a SC calcium current, and *I_SC_K_* is a SC potassium current. Each of the SC currents were fitted to the following equation:

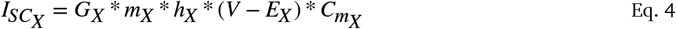

where *X* is Na, Ca, or K, G is a conductance, *m* is an activation coefficient, *h* is an inactivation coefficient, and *E* is a Nernst potential. The CM resting potential and action potential duration at 90% repolarization (APD90) were measured for *N_SC_* ranging from 0-5.

**Supplemental Figure 1.**
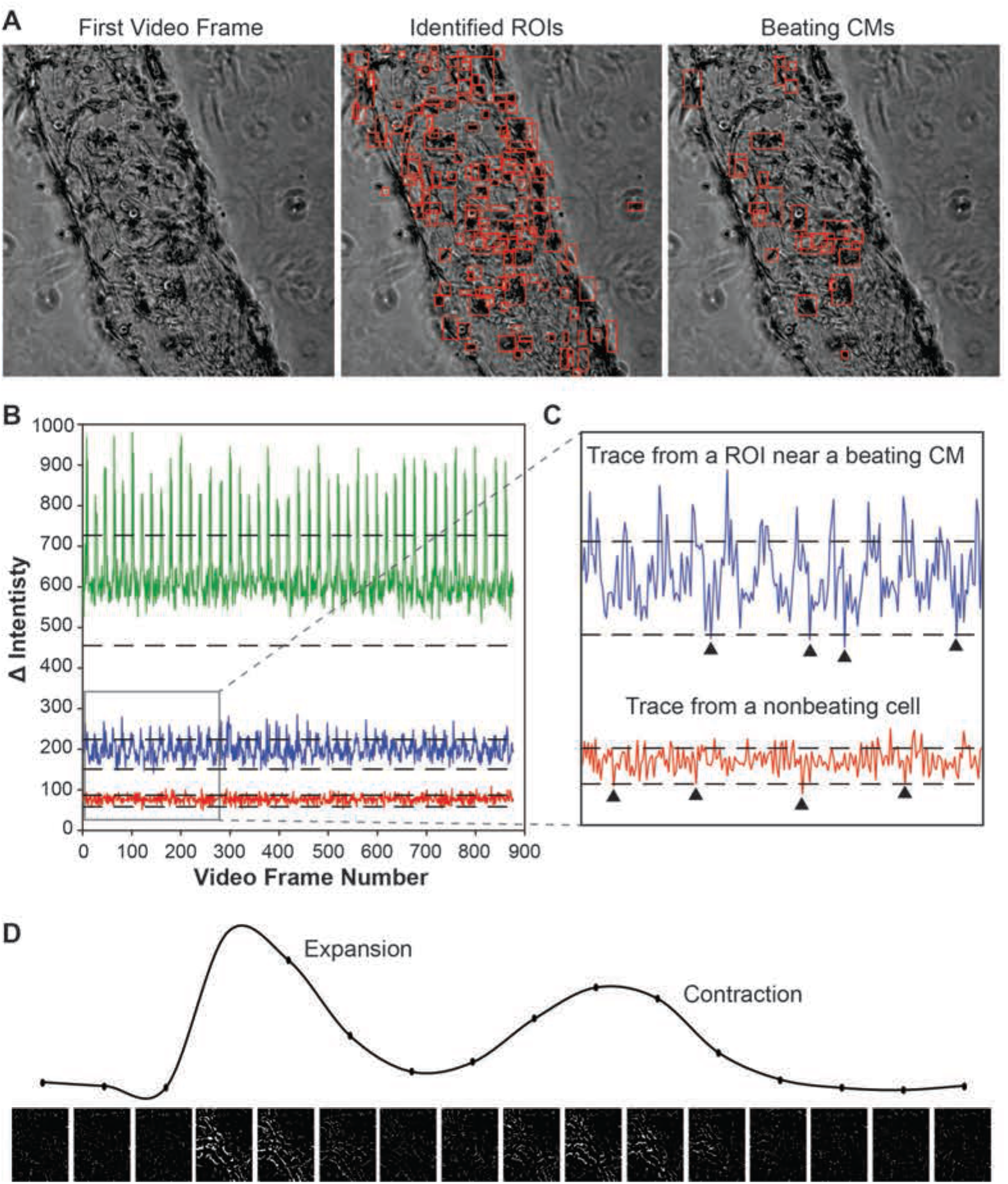
Cardiac beating algorithm inclusions criteria. **(A)** Representative phase contrast images of cardiac μTissues with all identified ROIs boxed in red and those identified to be beating CMs. **(B)** Representative summation of frame-to-frame differences from a beating CM (green), a nonbeating cell near a CM (blue), and a nonbeating cell (red). **(C)** Zoomed in profiles of the blue and red traces, denoted by boxed region in (B), showing peaks below the negative threshold value which excludes those ROIs from being identified as a beating CM. **(D)** Plot of the summation of frame-to-frame differences (computer generated images below) showing the expansion and contraction of CMs.

**Supplemental Figure 2.**
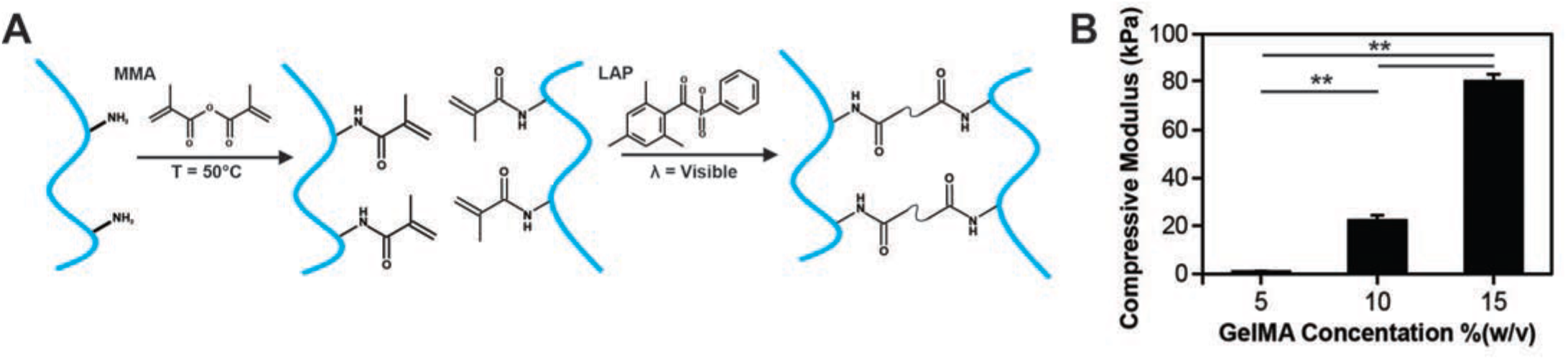
Material characterization. **(A)** Proposed chemical synthesis of visible light crosslinked GelMA hydrogels. **(B)** Compressive modulus of 5, 10, and 15 %(w/v) GelMA hydrogels. (** p < 0.05)

**Supplemental Figure 3.**
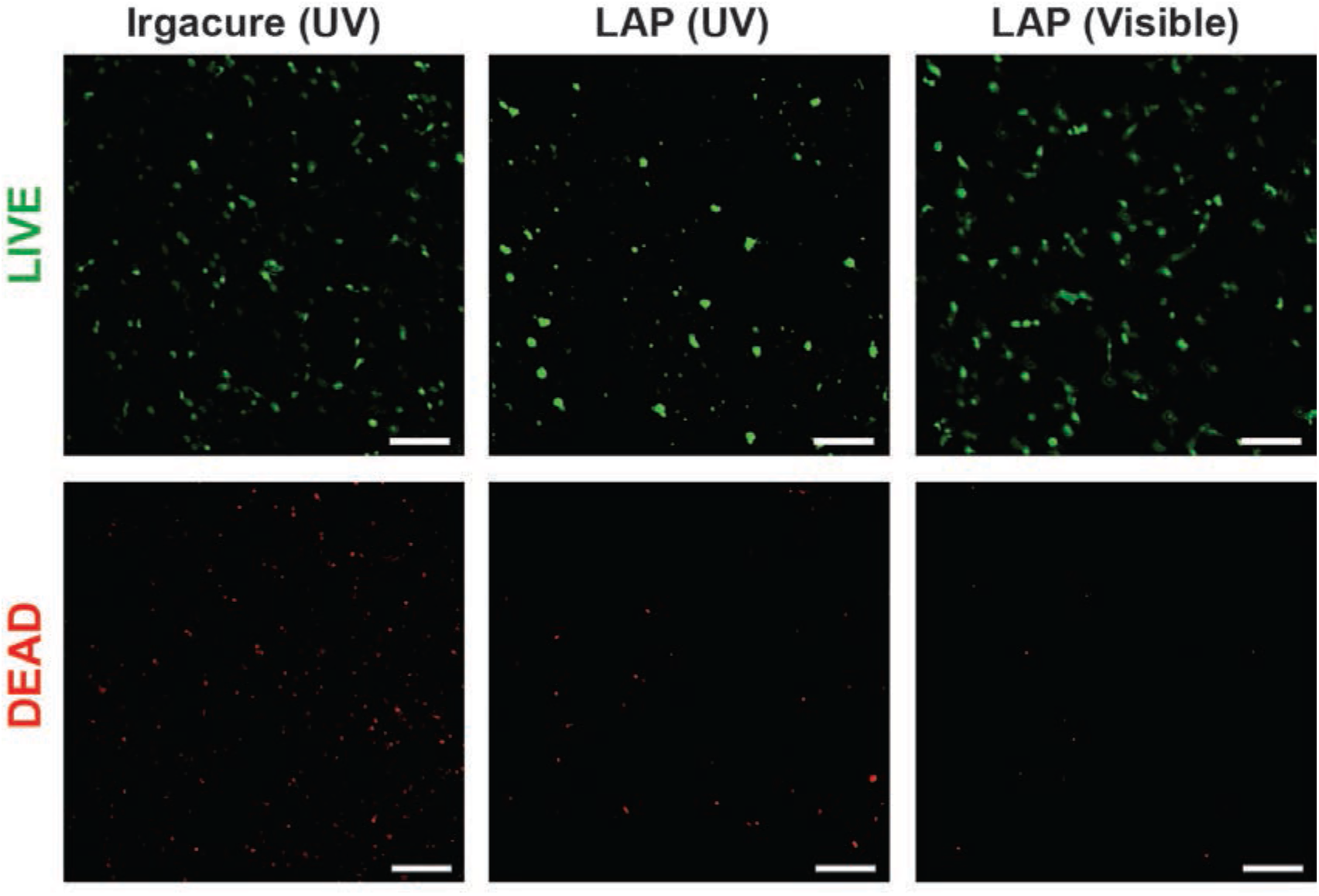
Cell viability. Representative live (green)/dead (red) images of cardiac cells encapsulated in GelMA hydrogels crosslinked with UV or visible light in the presence of Irgacure^®^ or LAP (scale = 200 μm).

**Supplemental Figure 4.**
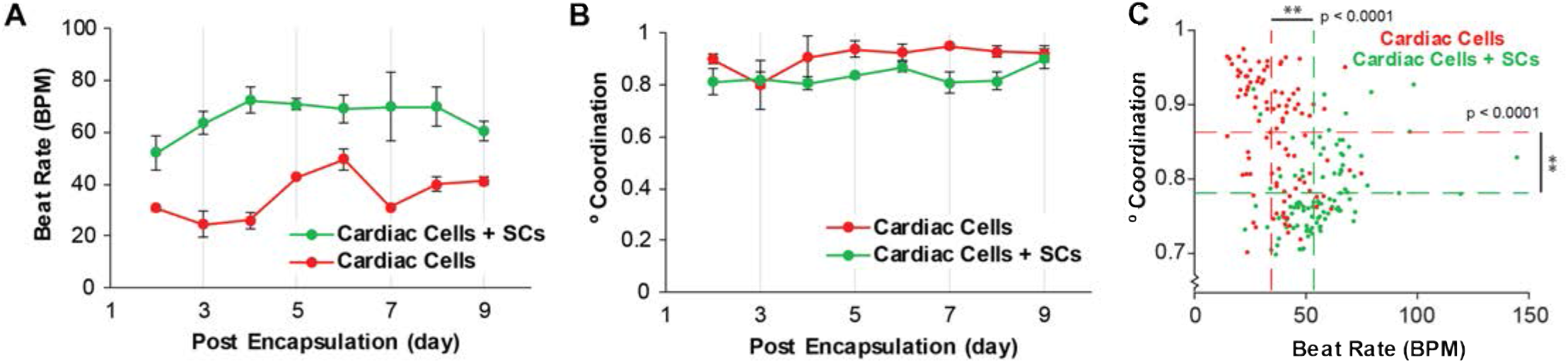
SCs role in altering cardiac output in 2D. **(A)** Quantification of beat rate and **(B)** beating synchrony for cardiac cells culture in 2D over time showing that the inclusions of SCs leads to an increase and beat rate and decrease if coordinated contracts. **(C)** Multi-way ANOVA showing statistical differences between beat rate and coordination for cardiac cells cultures (in both 2D/3D) with (green) and without (red) the addition of exogenous SC. (** p < 0.05)

**Supplemental Figure 5.**
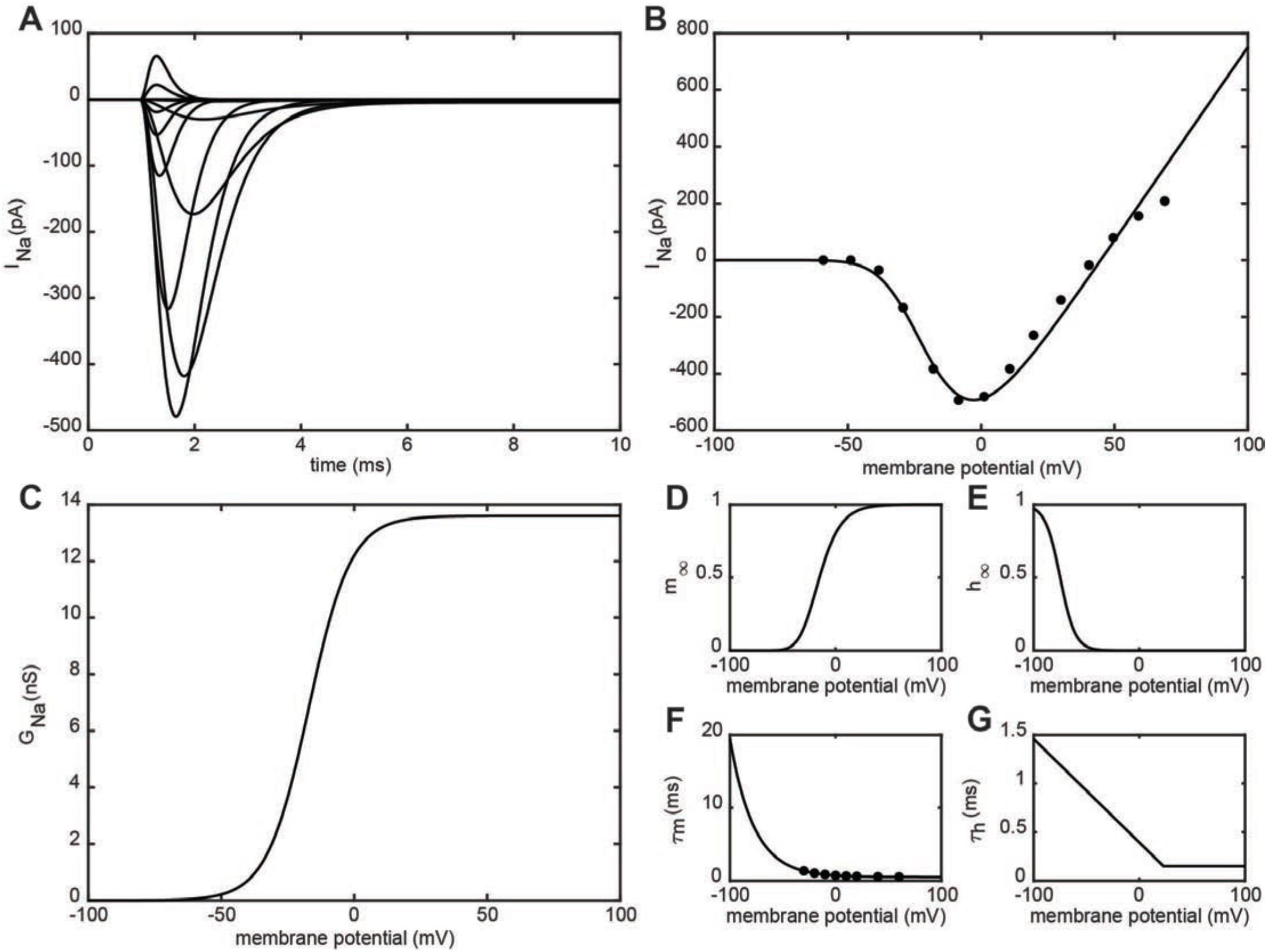
*In silico* modeling of a Schwann cell sodium channel. **(A)** Computational voltage clamp stimulation for a SC sodium channel from −60 mV to 60 mV with 10 mV steps and a resting potential of −70 mV. **(B)** Plot for max current vs potential plot curve fit by the following equation: I_Na_ = 13.6 * m_∞_^3^ * (1 - h_∞_ * (V - 44.8). **(C)** Plot of conductance at different membrane voltages given by the following equation: G_Na_ = 13.6 * 1/(1 + e^(−17.1 − V)/7.9^). **(D)** Plots of activation and **(E)** inactivation parameters given by the following equations: m_∞_ = 1/(1 + e^−(V + 32)/12.42^), and h_∞_ = 1/(1 + e^(V + 74.9)/7.08^). **(F)** Plots of the activation and **(G)** inactivation time constants curve fit and estimated by the following equations: **τ**_m_ = 0.2238 * e^−0.04441 * V^ + 0.5 * e^0.0001947 * V^, and **τ**_h_ = −(V - 37)/.094/1000, but for **τ**_h_ < 0.15, set **τ**_h_ = 0.15.

**Supplemental Figure 6.**
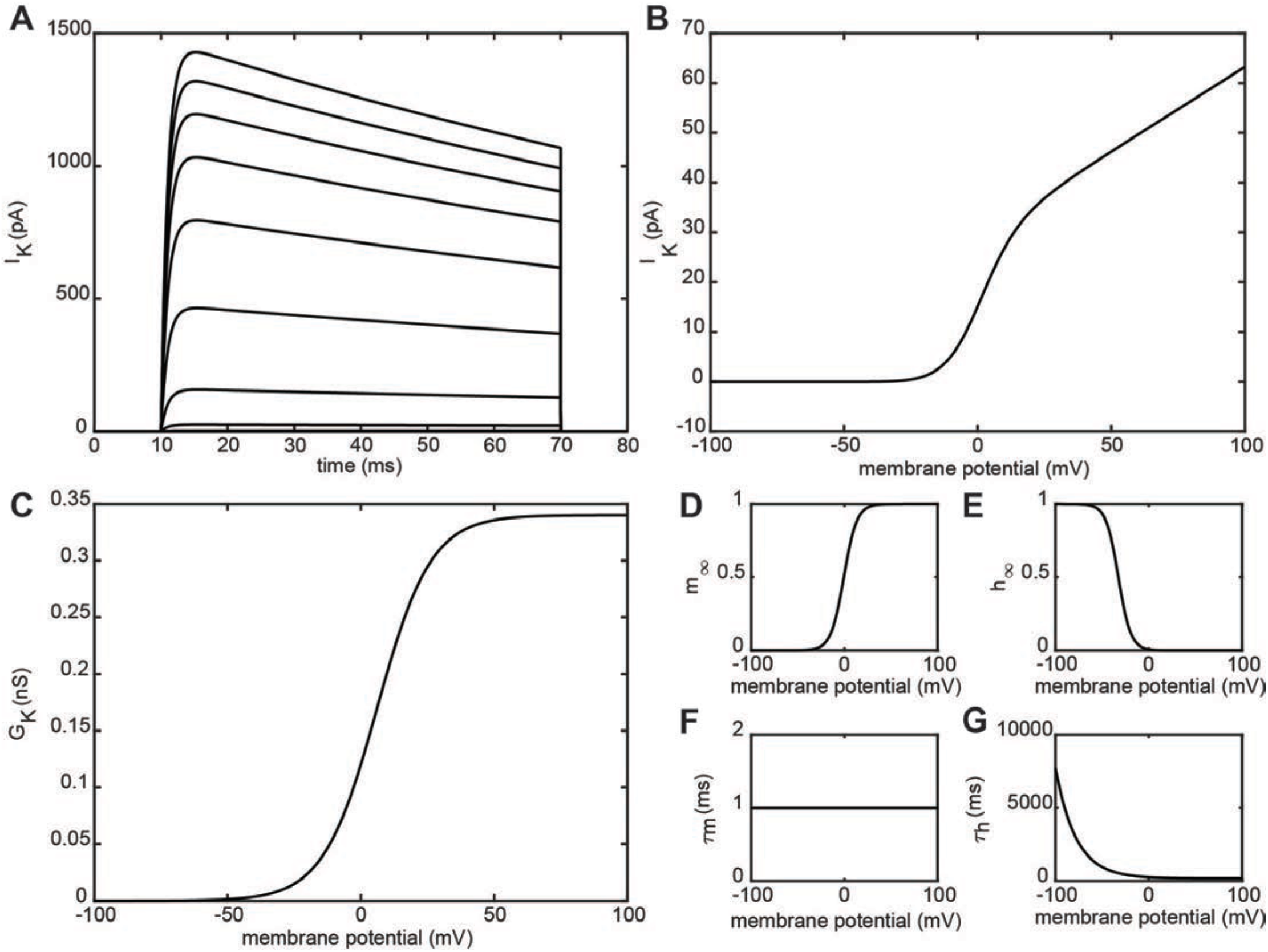
*In silico* modeling of a Schwann cell potassium channel. **(A)** Computational voltage clamp stimulation for a SC potassium channel from −60 mV to 60 mV with 10 mV steps and a resting potential of −70 mV. **(B)** Plot for max current vs potential plot modeled by the following equation: I_K_ = 0.34 * m_∞_ * (1 - h_∞_) * (V + 86). **(C)** Plot of conductance at different membrane voltages given by the following equation: G_Na_ = 0.34 * 1/(1 + e^(6.1 - V)/10.1^). **(D)** Plots of activation and **(E)** inactivation parameters given by the following equations: m_∞_ = 1/(1 + e^−(V + 0.6)/6.9^), and h_∞_ = 1/(1 + e^(V + 32.6)/6.6^). **(F)** Plots of the activation and **(G)** inactivation time constants estimated by the following equations: **τ**_m_ = 1, and **τ**_h_ = 100 * (1.8 + 1/(1.25 * e^V/22^)).

**Supplemental Figure 7.**
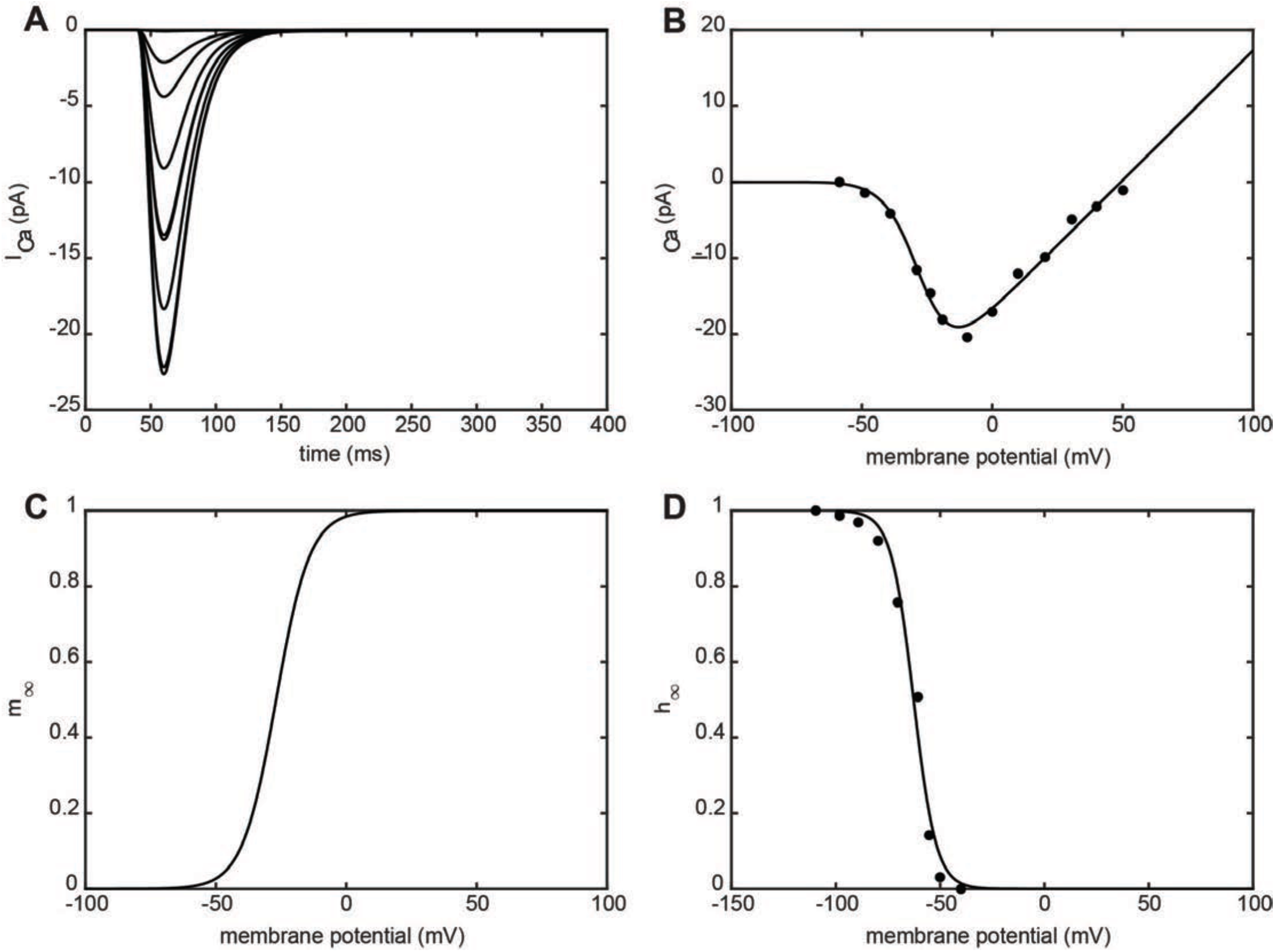
*In silico* modeling of a Schwann cell calcium channel. **(A)** Computational voltage clamp stimulation for a SC calcium channel from −40 mV to 40 mV with 10 mV steps and a resting potential of −70 mV. **(B)** Plot for max current vs potential plot curve fit by the following equation: I_K_ = 0.3419 * m_∞_ * (1 - h_∞_) * (V − 49.3). **(C)** Plots of activation and **(D)** inactivation parameters given and curve fit by the following equations: m_∞_ = 1/(1 + e^−(V + 26.9)/6.47^), and h_∞_ = 1/(1 + e^(V + 62.7)/5.41^). Activation and inactivation time constants were estimated to be constant values: **τ**_m_ = 15, and **τ**_h_ = 15.

**Supplemental Figure 8.**
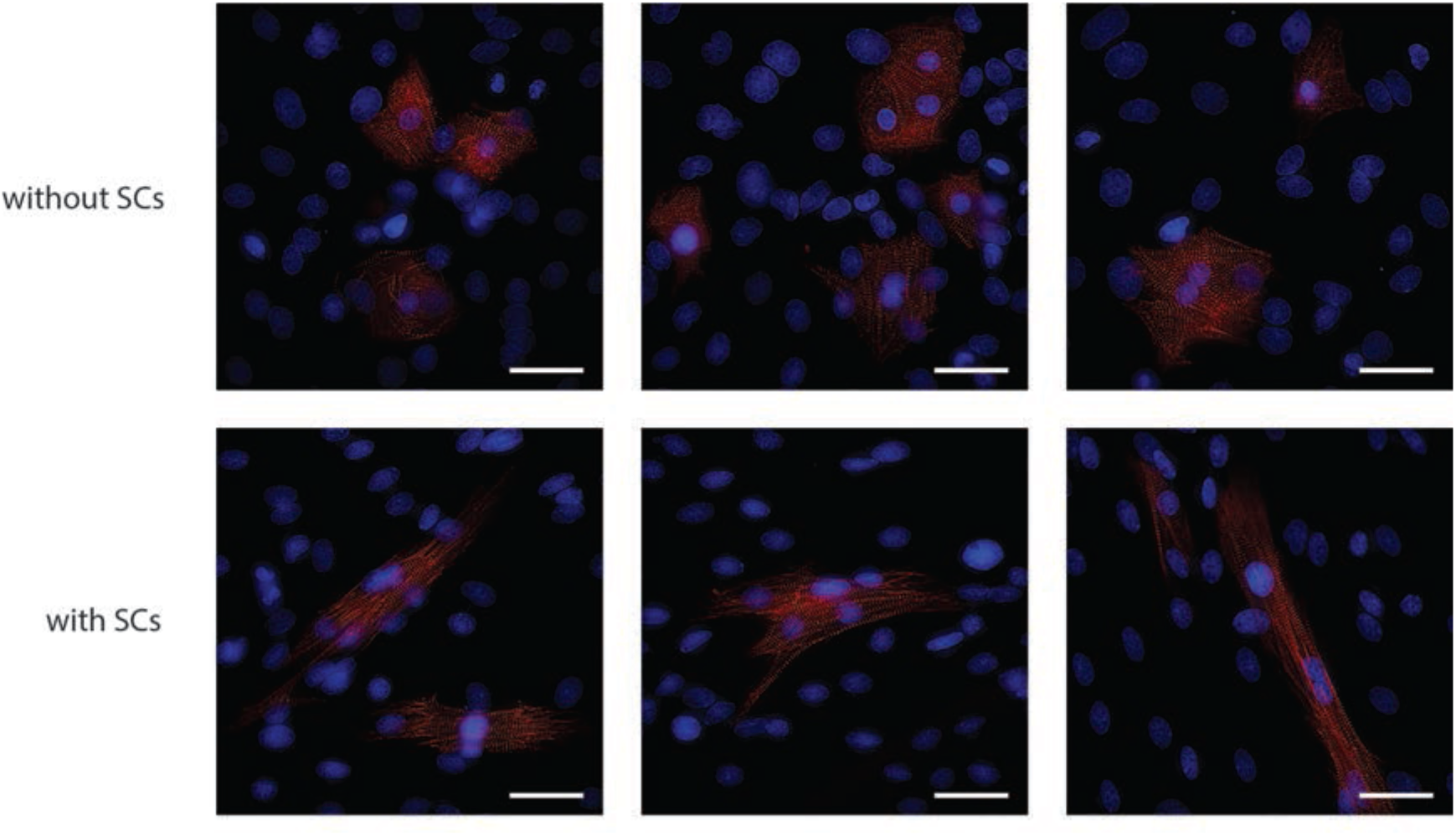
SCs role on CM morphology. Representative images (red: sarcomeric α-actinin, blue: DAPI) showing increased binucleation with the inclusion of SCs (scale = 40 μm).

**Supplemental Figure 9.**
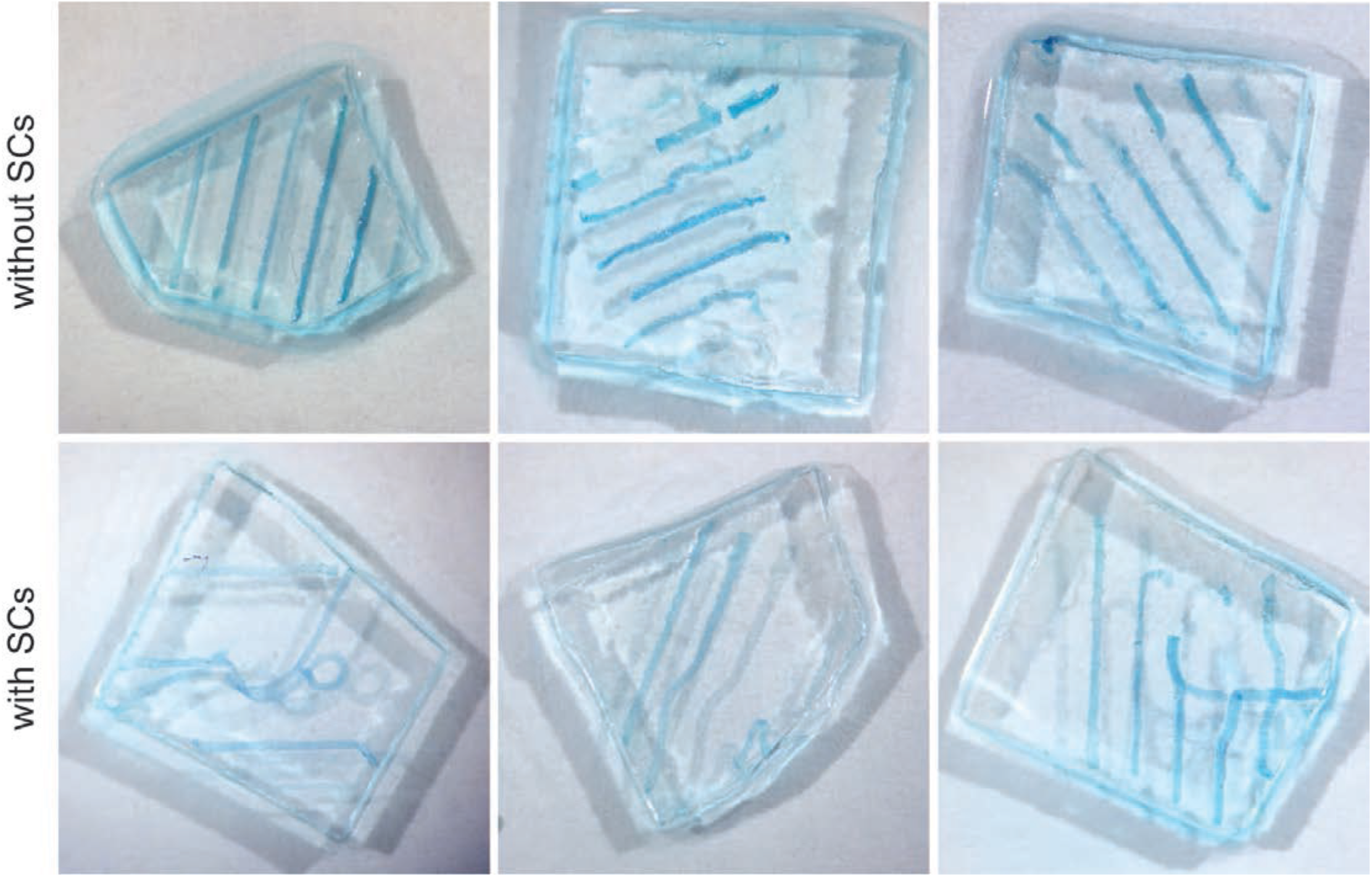
SCs impact on material degradation. Representative photographs of patterned hydrogels on day 9 post encapsulation showing the inclusion of SCs in 3D cardiac μTissues leads to delamination of the hydrogel from the glass. Samples were incubated in food coloring to improve contrast.

## References

1 Björnson, E., Borén, J. & Mardinoglu, A. Personalized Cardiovascular Disease Prediction and Treatment—A Review of Existing Strategies and Novel Systems Medicine Tools. Frontiers in Physiology 7, 2, doi:10.3389/fphys.2016.00002 (2016).

2 Malliaras K. & Marbán E. Cardiac cell therapy: where we’ve been, where we are, and where we should be headed. British Medical Bulletin 98, 161–185, doi:10.1093/bmb/ldr018 (2011).

3 Zhou P. & Pu, W. T. Recounting Cardiac Cellular Composition. Circ. Res. 118, 368–370, doi:10.1161/CIRCRESAHA.116.308139 (2016).

4 Pinto A. R. et al. Revisiting Cardiac Cellular Composition. Circ. Res. 118, 400–409, doi:10.1161/CIRCRESAHA.115.307778 (2016).

5 Popescu L. M. et al. Telocytes and putative stem cells in ageing human heart. Journal of Cellular and Molecular Medicine 19, 31–45, doi:10.1111/jcmm.12509 (2015).

6 Nandi S. S. & Mishra P. K. Harnessing fetal and adult genetic reprograming for therapy of heart disease. Journal of nature and science 1 (2015).

7 Gherghiceanu M. & Popescu L. M. Cardiac telocytes — their junctions and functional implications. Cell and Tissue Research 348, 265–279, doi:10.1007/s00441-012-1333-8 (2012).

8 Xin, M., Olson, E. N. & Bassel-Duby, R. Mending broken hearts: cardiac development as a basis for adult heart regeneration and repair. Nat. Rev. Mol. Cell Biol. 14, doi:10.1038/nrm3619 (2013).

9 Armour A. J. Intrinsic Cardiac Neurons. Journal of Cardiovascular Electrophysiology 2, 331–341, doi:10.1111/j.1540-8167.1991.tb01330.x (1991).

10 Iseoka H. et al. Pivotal Role of Non-cardiomyocytes in Electromechanical and Therapeutic Potential of Induced Pluripotent Stem Cell-Derived Engineered Cardiac Tissue. Tissue Eng Part A 24, 287–300, doi:10.1089/ten.TEA.2016.0535 (2018).

11 Sachse, F. B., Moreno, A. P. & Abildskov, J. A. Electrophysiological Modeling of Fibroblasts and their Interaction with Myocytes. Ann. Biomed. Eng. 36, 41–56, doi:10.1007/s10439-007-9405-8 (2008).

12 Hulsmans M. et al. Macrophages Facilitate Electrical Conduction in the Heart. Cell 169, 510, doi:10.1016/j.cell.2017.03.050 (2017).

13 Sheng J. et al. Electrophysiology of human cardiac atrial and ventricular telocytes. Journal of Cellular and Molecular Medicine 18, 355–362, doi:10.1111/jcmm.12240 (2014).

14 Mayourian, J., Savizky, R. M., Sobie, E. A. & Costa, K. D. Modeling Electrophysiological Coupling and Fusion between Human Mesenchymal Stem Cells and Cardiomyocytes. PLoS Comput. Biol. 12, doi:10.1371/journal.pcbi.1005014 (2016).

15 Natarajan A. et al. Patterned cardiomyocytes on microelectrode arrays as a functional, high information content drug screening platform. Biomaterials 32, 4267–4274, doi:10.1016/j.biomaterials.2010.12.022 (2011).

16 Zuppinger C. 3D culture for cardiac cells. Biochimica et Biophysica Acta (BBA)-Molecular Cell Research 1863, 1873–1881, doi:10.1016/j.bbamcr.2015.11.036 (2016).

17 Li, X., Valadez, A. V., Zuo, P. & Nie, Z. Microfluidic 3D cell culture: potential application for tissue-based bioassays. Bioanalysis 4, 1509–1525, doi:10.4155/bio.12.133 (2012).

18 Tibbitt M. W. & Anseth K. S. Hydrogels as extracellular matrix mimics for 3D cell culture. Biotechnol. Bioeng. 103, 655–663, doi:10.1002/bit.22361 (2009).

19 Mandrycky, C., Wang, Z., Kim, K. & Kim, D.-H. 3D bioprinting for engineering complex tissues. Biotechnol. Adv. 34, 422–434, doi:10.1016/j.biotechadv.2015.12.011 (2016).

20 Yue K. et al. Synthesis, properties, and biomedical applications of gelatin methacryloyl (GelMA) hydrogels. Biomaterials 73, 254–271, doi:10.1016/j.biomaterials.2015.08.045 (2015).

21 DeForest, C. A., Polizzotti, B. D. & Anseth, K. S. Sequential click reactions for synthesizing and patterning three-dimensional cell microenvironments. Nat. Mater. 8, doi:10.1038/nmat2473 (2009).

22 Noshadi I. et al. In vitro and in vivo analysis of visible light crosslinkable gelatin methacryloyl (GelMA) hydrogels. Biomaterials Science 5, 2093–2105, doi:10.1039/C7BM00110J (2017).

23 Nichol J. W. et al. Cell-laden microengineered gelatin methacrylate hydrogels. Biomaterials 31, 5536–5544, doi:10.1016/j.biomaterials.2010.03.064 (2010).

24 Noshadi I. et al. Engineering Biodegradable and Biocompatible Bio-ionic Liquid Conjugated Hydrogels with Tunable Conductivity and Mechanical Properties. Scientific Reports 7, 4345, doi:10.1038/s41598-017-04280-w (2017).

25 Koppes A. N. et al. Electrical stimulation of schwann cells promotes sustained increases in neurite outgrowth. Tissue Eng Part A 20, 494–506, doi:10.1089/ten.TEA.2013.0012 (2014).

26 Kreuz, T., Mulansky, M. & Bozanic, N. SPIKY: a graphical user interface for monitoring spike train synchrony. J Neurophysiol 113, 3432–3445, doi:10.1152/jn.00848.2014 (2015).

27 Tandon N. et al. Electrical stimulation systems for cardiac tissue engineering. Nat. Protoc. 4, 155–173, doi:10.1038/nprot.2008.183 (2009).

28 Baker M. D. & Ritchie J. M. Characteristics of type I and type II K^+^ channels in rabbit cultured Schwann cells. The Journal of physiology 490 (Pt 1), 79–95, doi:10.1113/jphysiol.1996.sp021128 (1996).

29 Amédée, T., Ellie, E., Dupouy, B. & Vincent, J. D. Voltage–dependent calcium and potassium channels in Schwann cells cultured from dorsal root ganglia of the mouse. The Journal of Physiology 441, 35–56, doi:10.1113/jphysiol.1991.sp018737 (1991).

30 Howe J. R. & Ritchie J. M. Sodium currents in Schwann cells from myelinated and non-myelinated nerves of neonatal and adult rabbits. The Journal of physiology 425, 169–210 (1990).

31 Bhana B. et al. Influence of substrate stiffness on the phenotype of heart cells. Biotechnol Bioeng 105, 1148–1160, doi:10.1002/bit.22647 (2010).

32 Huebsch N. et al. Automated Video-Based Analysis of Contractility and Calcium Flux in Human-Induced Pluripotent Stem-Derived Cardiomyocytes Cultured over Different Spatial Scales. Tissue Engineering Part C: Methods 21, 467–479, doi:10.1089/ten.tec.2014.0283 (2015).

33 Saini, H., Navaei, A., Van Putten, A. & Nikkhah, M. 3D cardiac microtissues encapsulated with the co-culture of cardiomyocytes and cardiac fibroblasts. Adv Healthc Mater 4, 1961–1971, doi:10.1002/adhm.201500331 (2015).

34 Annabi N. et al. Highly Elastic Micropatterned Hydrogel for Engineering Functional Cardiac Tissue. Adv Funct Mater 23, 4950–4959, doi:10.1002/adfm.201300570 (2013).

35 Maoz B. M. et al. Organs-on-Chips with combined multi-electrode array and transepithelial electrical resistance measurement capabilities. Lab Chip 17, 2294–2302, doi:10.1039/C7LC00412E (2017).

36 Zhang H. et al. Alteration of parasympathetic/sympathetic ratio in the infarcted myocardium after Schwann cell transplantation modified electrophysiological function of heart: a novel antiarrhythmic therapy. Circulation 122, S193–200, doi:10.1161/CIRCULATI0NAHA.109.922740 (2010).

37 Wang Y. et al. Transplantation of microencapsulated Schwann cells and mesenchymal stem cells augment angiogenesis and improve heart function. Mol Cell Biochem 366, 139–147, doi:10.1007/s11010-012-1291-1 (2012).

38 Blaeser A. et al. Controlling Shear Stress in 3D Bioprinting is a Key Factor to Balance Printing Resolution and Stem Cell Integrity. Advanced Healthcare Materials 5, 326–333, doi:10.1002/adhm.201500677 (2016).

39 Annabi N. et al. Engineering a sprayable and elastic hydrogel adhesive with antimicrobial properties for wound healing. Biomaterials 139, 229–243, doi:10.1016/j.biomaterials.2017.05.011 (2017).

40 Fairbanks, B. D., Schwartz, M. P., Bowman, C. N. & Anseth, K. S. Photoinitiated polymerization of PEG-diacrylate with lithium phenyl-2,4,6-trimethylbenzoylphosphinate: polymerization rate and cytocompatibility. Biomaterials 30, 6702–6707, doi:10.1016/j.biomaterials.2009.08.055 (2009).

41 Maxwell S. R. & Lip G. Y. Free radicals and antioxidants in cardiovascular disease. British journal of clinical pharmacology 44, 307–317 (1997).

42 Morimoto, Y., Kato-Negishi, M., Onoe, H. & Takeuchi, S. Three-dimensional neuron-muscle constructs with neuromuscular junctions. Biomaterials 34, 9413–9419, doi:10.1016/j.biomaterials.2013.08.062 (2013).

43 Uzel S. G. et al. Microfluidic device for the formation of optically excitable, three-dimensional, compartmentalized motor units. Sci Adv 2, e1501429, doi:10.1126/sciadv.1501429 (2016).

44 Boudou T. et al. A Microfabricated Platform to Measure and Manipulate the Mechanics of Engineered Cardiac Microtissues. Tissue Eng. Part A 18, 910–919, doi:10.1089/ten.tea.2011.0341 (2012).

45 Armati P. J. & Mathey E. K. An update on Schwann cell biology–immunomodulation, neural regulation and other surprises. J. Neurol. Sci. 333, 68–72, doi:10.1016/j.jns.2013.01.018 (2013).

46 Zhang H. et al. Alteration of Parasympathetic/Sympathetic Ratio in the Infarcted Myocardium After Schwann Cell Transplantation Modified Electrophysiological Function of Heart A Novel Antiarrhythmic Therapy. Circulation 122, S193–S200, doi:10.1161/CIRCULATIONAHA.109.922740 (2010).

47 Annabi N. et al. Highly Elastic and Conductive Human-Based Protein Hybrid Hydrogels. Adv Mater 28, 40–49, doi:10.1002/adma.201503255 (2016).

48 Shin S. et al. Reduced Graphene Oxide-GelMA Hybrid Hydrogels as Scaffolds for Cardiac Tissue Engineering. Small 12, 3677–3689, doi:10.1002/smll.201600178 (2016).

49 Denning C. et at. Cardiomyocytes from human pluripotent stem cells: From laboratory curiosity to industrial biomedical platform. BBA 1863, 1728–1748, doi:10.1016/j.bbamcr.2015.10.014 (2016).

50 Scuderi G. J. & Butcher J. Naturally Engineered Maturation of Cardiomyocytes. Front Cell Dev Biol 5, 50, doi:10.3389/fcell.2017.00050 (2017).

51 Zebrowski D. C. et al. Cardiac injury of the newborn mammalian heart accelerates cardiomyocyte terminal differentiation. Sci Rep 7, 8362, doi:10.1038/s41598-017-08947-2 (2017).

52 Hoffman-Kim, D., Mitchel, J. A. & Bellamkonda, R. V. Topography, cell response, and nerve regeneration. Annu Rev Biomed Eng 12, 203–231, doi:10.1146/annurev-bioeng-070909-105351 (2010).

53 Seggio, A. M., Narayanaswamy, A., Roysam, B. & Thompson, D. M. Self-aligned Schwann cell monolayers demonstrate an inherent ability to direct neurite outgrowth. J Neural Eng. 7, 46001, doi:10.1088/1741-2560/7/4/046001 (2010).

54 Lindsey, S. E., Butcher, J. T. & Yalcin, H. C. Mechanical regulation of cardiac development. Front Physiol 5, 318, doi:10.3389/fphys.2014.00318 (2014).

55 Liu H. et al. Matrix metalloproteinase inhibition enhances the rate of nerve regeneration in vivo by promoting dedifferentiation and mitosis of supporting schwann cells. J Neuropathol Exp Neurol 69, 386–395, doi:10.1097/NEN.0b013e3181d68d12 (2010).

56 Parrinello S. et al. EphB signaling directs peripheral nerve regeneration through Sox2-dependent Schwann cell sorting. Cell 143, 145–155, doi:10.1016/j.cell.2010.08.039 (2010).

57 Vandooren, J., Van den Steen, P. E. & Opdenakker, G. Biochemistry and molecular biology of gelatinase B or matrix metalloproteinase-9 (MMP-9): the next decade. Crit Rev Biochem Mol Biol 48, 222–272, doi:10.3109/10409238.2013.770819 (2013).

58 Louch, W. E., Koivumäki, J. T. & Tavi, P. Calcium signalling in developing cardiomyocytes: implications for model systems and disease. The Journal of Physiology 593, 1047–1063, doi:10.1113/jphysiol.2014.274712 (2015).

59 Reist N. E. & Smith S. J. Neurally evoked calcium transients in terminal Schwann cells at the neuromuscular junction. Proc Natl Acad Sci U S A 89, 7625–7629 (1992).

60 Robert A. & Jirounek P. Uptake of potassium by nonmyelinating Schwann cells induced by axonal activity. J Neurophysiol 72, 2570–2579, doi:10.1152/jn.1994.72.6.2570 (1994).

61 Nicholson S. M. & Bruzzone R. Gap junctions: Getting the message through. Current Biology 7, doi:10.1016/S0960-9822(06)00169-2 (1997).

62 Brokamp, C., Todd, J., Montemagno, C. & Wendell, D. Electrophysiology of Single and Aggregate Cx43 Hemichannels. PLoS One 7, doi:10.1371/journal.pone.0047775 (2012).

63 Cimetta, E., Godier-Furnémont, A. & Vunjak-Novakovic, G. Bioengineering heart tissue for in vitro testing. Curr. Opin. Biotechnol. 24, 926–932, doi:10.1016/j.copbio.2013.07.002 (2013).

